# Connectome caricatures: removing large-amplitude co-activation patterns in resting-state fMRI emphasizes individual differences

**DOI:** 10.1101/2024.04.08.588578

**Authors:** Raimundo X. Rodriguez, Stephanie Noble, Chris C. Camp, Dustin Scheinost

## Abstract

High-amplitude co-activation patterns are sparsely present during resting-state fMRI but drive functional connectivity^1–5^. Further, they resemble task activation patterns and are well-studied^3,5–10^. However, little research has characterized the remaining majority of the resting-state signal. In this work, we introduced caricaturing—a method to project resting-state data to a subspace orthogonal to a manifold of co-activation patterns estimated from the task fMRI data. Projecting to this subspace removes linear combinations of these co-activation patterns from the resting-state data to create Caricatured connectomes. We used rich task data from the Human Connectome Project (HCP)^11^ and the UCLA Consortium for Neuropsychiatric Phenomics^12^ to construct a manifold of task co-activation patterns. Caricatured connectomes were created by projecting resting-state data from the HCP and the Yale Test-Retest^13^ datasets away from this manifold. Like caricatures, these connectomes emphasized individual differences by reducing between-individual similarity and increasing individual identification^14^. They also improved predictive modeling of brain-phenotype associations. As caricaturing removes group-relevant task variance, it is an initial attempt to remove task-like co-activations from rest. Therefore, our results suggest that there is a useful signal beyond the dominating co-activations that drive resting-state functional connectivity, which may better characterize the brain’s intrinsic functional architecture.

## 1. Introduction

Functional connectivity at rest is driven by bursts of short, spatially distributed co-activation events, which make up only a small fraction of the scan^1–5^. These co-activations increase the temporal correlation between functional networks and the nonstationarity of the resting-state signal^1,3^ and mimic task-induced activity patterns^3,5–10^. However, they may be problematic because they account for only a small fraction of the signal and occur sporadically, leading to low reliability^14,15^ and predictive utility^16^.

Relatedly, two theoretical frameworks exist to emphasize individual differences in functional connectivity^14^. The *spotlight* approach “[blurs] the irrelevant features while retaining and enriching relevant ones”. Task-induced changes in the fMRI signal achieve this goal by consistently modulating the co-activations across individuals^14,17^. Since task paradigms intentionally elicit these strong co-activations, they are more densely sampled during tasks, driving a comparatively stable connectivity pattern. As a result, task-based connectomes exhibit greater within- and between-individual similarity, reliability, and predictive utility than resting-state connectomes. These improvements have been observed across classic task paradigms^14–16,18,19^, as well as modern naturalistic and movie paradigms^20–22^. Methods which transform resting-state connectomes into task-based connectomes similarly improve reliability and prediction^23,24^.

The second framework—the *caricature* approach—exaggerates “the most prominent features of each individual”^14^. Like a caricature, individuals would become less similar to each other, reducing between-individual similarity and increasing reliability and predictive utility^14^. While no work has investigated how to implement this thought experiment, removing the co-activations in resting-state fMRI might formalize it. As task-induced co-activation patterns lie on a low-dimensional manifold common across participants^25^, principal component analysis (PCA) of task fMRI data can flexibly identify these patterns in an unsupervised manner. Projecting resting-state data onto a subspace orthogonal to this manifold would remove linear combinations of these co-activation patterns from the resting-state data. Importantly, as the co-activation patterns are defined at the group level, the group-relevant variance would be removed, reducing between-individual similarity as a caricature would. Additionally, caricaturing offers a method to study the remaining signal in resting-state data with the dominant, task-like co-activations removed, which may better characterize the brain’s intrinsic functional architecture.

To evaluate caricaturing, we compared the within- and between-individual similarity of the projected data (i.e., Caricatured connectomes) to standard resting-state connectomes in three datasets. We also compared the reliability and predictive utility of Caricatured connectomes. Caricatured connectomes behaved like caricatures, exhibiting lower between-individual similarity but higher multivariate reliability and predictive utility. This novel method enhances the individual differences in existing resting-state data and—given the ease of collecting and harmonizing resting-state data over task data—helps motivate future data collection. These results also suggest that there is an informative signal beyond the dominating co-activation patterns that drive resting-state functional connectivity.

## 2. Results

To implement caricaturing, we projected the resting-state fMRI time series away from a manifold of task co-activation patterns, estimated from the task-based time series (**Figure 1A**). First, group-level principal components (PCs) are calculated by temporally concatenating task-based time series across tasks and participants and performing principal component analysis (PCA). Next, a projection matrix is created to project an fMRI time point into a subspace orthogonal to the top PCs. Multiplying this projection matrix by each time point in a resting-state scan creates new time series without information from the top PCs. Finally, connectomes are made as usual by correlating time series pairs.

**Figure 1:**
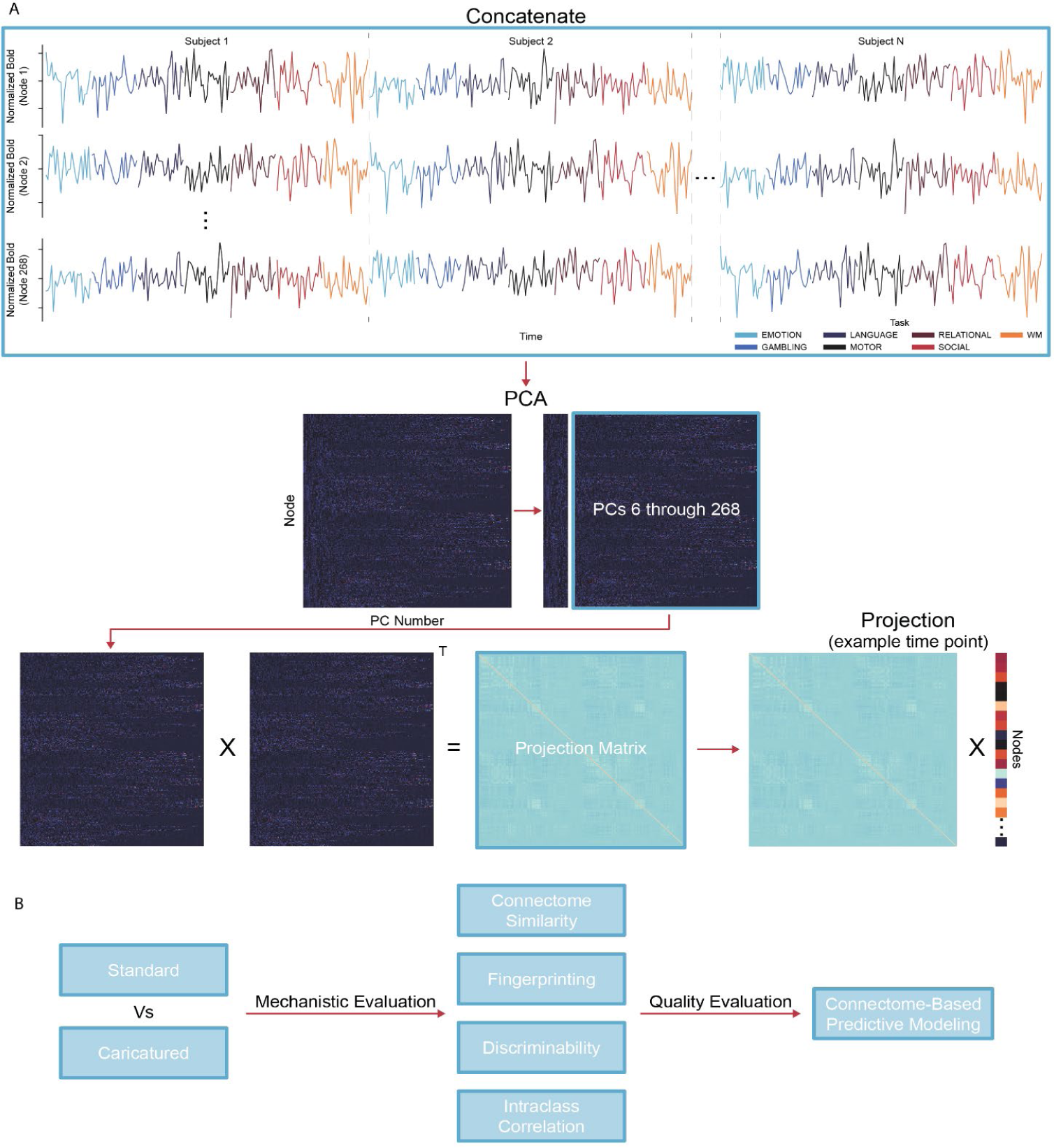
Caricaturing overview. **(A)** Caricaturing has two parts. The first is to define a task manifold from group-level task fMRI. We concatenate all task scans for individuals temporally and perform principal component analysis (PCA). Each principal component (PC) is a common co-activation pattern across tasks. The second part is to project resting-state data away from this manifold. We create a matrix of PCs excluding the top PCs (e.g., the first five, as in this work). This matrix is multiplied by its transpose (denoted as “T”) to obtain the projection matrix. Next, we multiply the projection matrix and each time point from a resting-state scan, orthogonalizing them to the task manifold. Caricatured connectomes are created by correlating these orthogonalized time series. **(B)** In several downstream analyses, we compare Caricatured connectomes to Standard connectomes (i.e., those created using standard resting-state fMRI data). First, we observed how the method mechanistically changes the data by investigating within- and between-individual similarity and multivariate (fingerprinting and discriminability) and univariate (edge-level intraclass correlation) reliability. Next, we used Connectome-Based Predictive Modeling to investigate if mechanistic changes in Caricatured connectomes resulted in stronger brain-behavior associations.

We used the Human Connectome Project (HCP)^11^, UCLA Consortium for Neuropsychiatric Phenomics (CNP)^12^, and the Yale Test-Retest (TRT) datasets, the latter of which comprises the publicly available TRTI^13^ and the private TRTII. The HCP and CNP datasets have rich task data and were used to create the manifold of task co-activations and associated projection matrices, independently. Using two datasets to create different manifolds highlights that caricaturing is general to the tasks and datasets used. Resting-state data from the HCP and TRT datasets were projected away from the manifold, creating ‘Caricatured’ connectomes. ‘Caricatured_HCP_’ or ‘Caricatured_CNP_’ indicates which dataset–the HCP or CNP– was used to generate the manifold. When using only the HCP to create Caricatured connectomes, we used a validated subsampling procedure (**SI Figures 1-3**) to avoid data leakage. **SI Figure 4** highlights the diminished correlation structure of the Caricatured connectomes compared to standard connectomes.

Caricatured connectomes from HCP and TRT were evaluated and compared to ‘Standard’ connectomes (i.e., those created using standard resting-state fMRI data) in multiple downstream analyses (**Figure 1B**). First, we characterized within-and between-participant similarity in the HCP and TRT datasets. Second, we analyzed multivariate reliability via fingerprinting and discriminability and univariate reliability via intra-class correlation (ICC) in the HCP and TRT datasets. Multivariate reliability reflects the stability of multidimensional data, such as whole-brain patterns, while univariate reliability reflects the reliability of each measurement individually. For the similarity and reliability analyses, we truncated the HCP time series to 176 frames before constructing the connectomes as was done in Finn et al., 2017^14^ to avoid a potential ceiling effect since longer scan duration improves connectome stability^26–29^ and fingerprinting accuracy^14^. Third, we performed connectome-based predictive modeling (CPM) in the HCP dataset for age, IQ, and sex, comparing prediction performance between Caricatured and Standard connectomes. Models for age and IQ used ridge regression; models for sex used support vector machines (SVM). Models were trained and tested using 1000 iterations of 10-fold cross-validation.

### 2.1 Group-level task-based PCA replication and validation

In the HCP, we replicated previous results^25^ to verify that task-driven activity lies on a low-dimensional manifold. PCA was performed on the concatenated task data from every participant in the HCP dataset. We then correlated the top five PC time series with the task block regressors. In line with previous results, the first PC time series was strongly correlated with the full task block regressor, created by combining the regressors across all tasks (mean r=0.387; **SI Figure 1A**). Additionally, the fifth PC time series was correlated with the absolute value of the derivative of the full task block regressors (mean r=0.113). The remaining PC time series differentially correlated with the task regressors (**SI Figure 1B**).

### 2.2 Caricatured connectomes decrease within- and between-individual similarity

We investigated within- and between-individual similarity for Caricatured and Standard connectomes as a caricature should decrease between-individual similarity. As expected in the HCP, Caricatured_HCP_ connectomes showed 53% lower between-individual similarity (p’s<0.008; Bonferroni corrected; Figure 2A). However, these connectomes also showed 42% less within-individual similarity (p’s<0.008; Bonferroni corrected; Figure 2A). Likewise, Caricatured_CNP_ connectomes exhibited a 39% decrease in within-individual similarity and a 48% decrease in between-individual similarity (p’s<0.0001; Bonferroni corrected; Figure 2B). Lastly, in the TRT dataset, Caricatured connectomes followed suit (Figure 2C) with significantly decreased within- (25%) and between-individual (41%) similarity (p’s<0.0001; Bonferroni corrected). Notably, these changes contrast with spotlighting, which increases both within- and between-individual similarity^14^.

**Figure 2:**
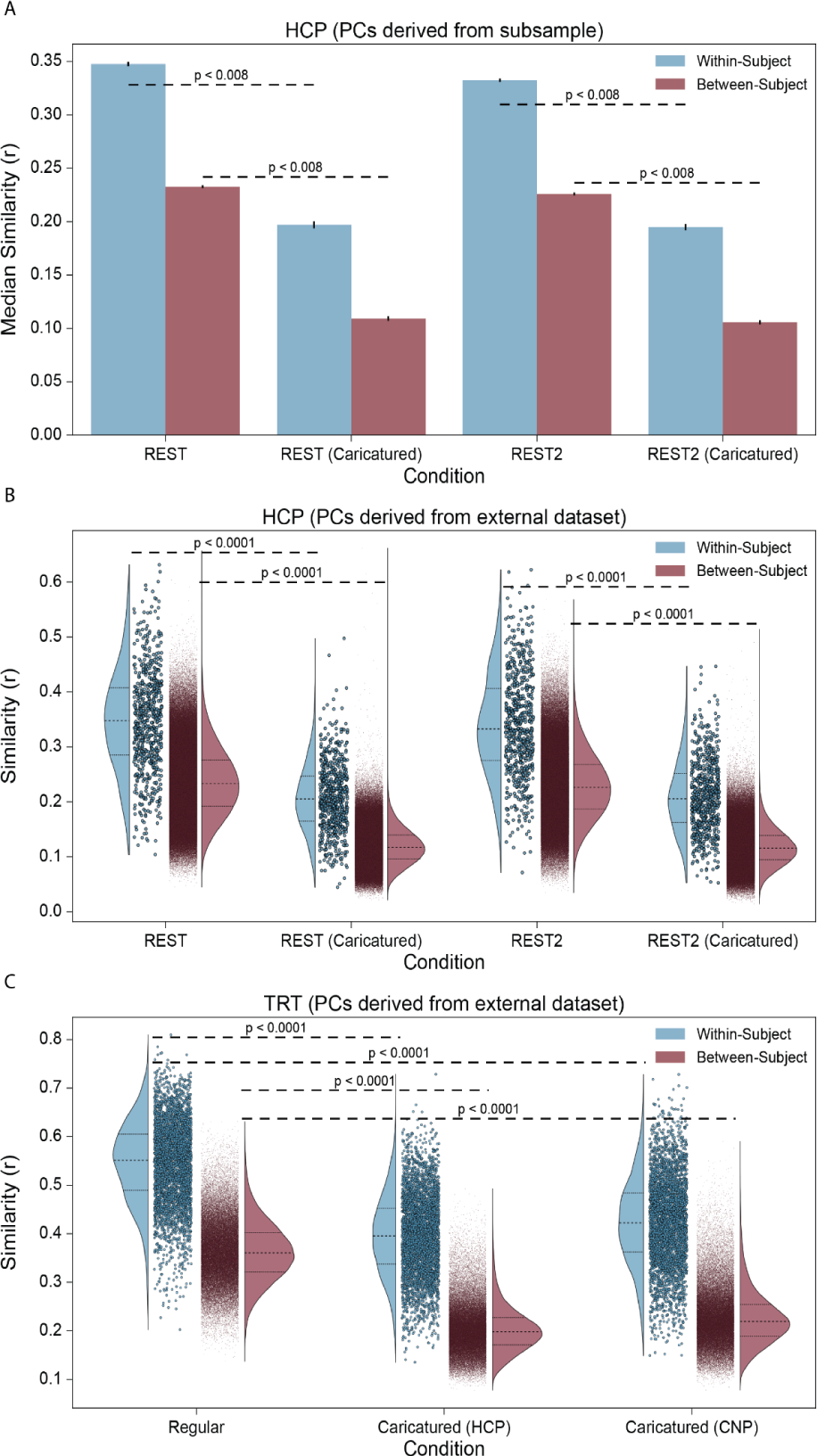
Connectome similarity. **(A)** Using 1000 iterations of the PC subsample procedure, the similarity between LR and RL phase-encoded scan pairs was calculated in the HCP dataset within each scan condition. Each iteration obtained within-individual and between-individual similarity distributions, and the median was taken for each distribution. Blue bars show the average median within-individual similarity, with error bars showing the standard deviation. Red bars show the same for the median between-individual similarity. **(B)** The same analysis was performed, instead using the CNP dataset to construct the PCs. Only one iteration was performed, with all participants used. The full within-individual similarity distribution (blue) and between-individual similarity distribution (red) are shown for each condition. **(C)** The analysis was performed in the TRT dataset, using either the HCP or CNP dataset to construct the PCs for projection. The full within-individual similarity distribution (blue) and between-individual similarity distribution (red) are shown for each condition. P-values are shown for all relevant comparisons; they are denoted as “less than” a particular value when the resolution of the test could not be discerned below that value. We chose 0.0001 as the lowest bound above which to report p-values.

### 2.3 Caricaturing improves multivariate reliability

We investigated multivariate reliability to determine whether changes in connectome similarity had a wider effect on the data. Reliability is broadly a function of within- and between-individual similarity, where a higher ration between the two is associated with higher reliability. In the HCP dataset, Caricatured_HCP_ connectomes exhibited significantly better fingerprinting than Standard connectomes, increasing accuracy by 41% (p’s<0.004; Bonferroni corrected; Figure 3A). Results were similar when using Caricatured_CNP_ connectomes, with a 42% average increase in accuracy (p’s<0.004; Bonferroni corrected; **SI Table 1**). In the TRT dataset, we also assessed the perfect separability rate (PSR)^13^, an extension of fingerprinting for datasets with more than two scan sessions per participant. Caricatured connectomes improved PSR by 508% (p’s<0.004; Bonferroni corrected; **SI Table 2**). Using the HCP, Caricatured_HCP_ and Caricatured_CNP_ connectomes increased discriminability on average by 4% compared to their Standard counterparts (p’s<0.004; Bonferroni corrected; Figure 3B; **SI Table 3**). Lastly, in the TRT dataset, discriminability was 1% higher for Caricatured_HCP_ and Caricatured_CNP_ connectomes than Standard connectomes (p_HCP_=0.28, p_CNP_=0.528; Bonferroni corrected; **SI Table 3**). Discriminability was near-perfect in the TRT dataset, leaving little room for large improvements. Likely, the small number of subjects inflates discriminability. While both within- and between-individual similarity decreased with Caricatured connectomes, between-individual similarity decreased more than within-individual similarity, increasing multivariate reliability.

**Figure 3:**
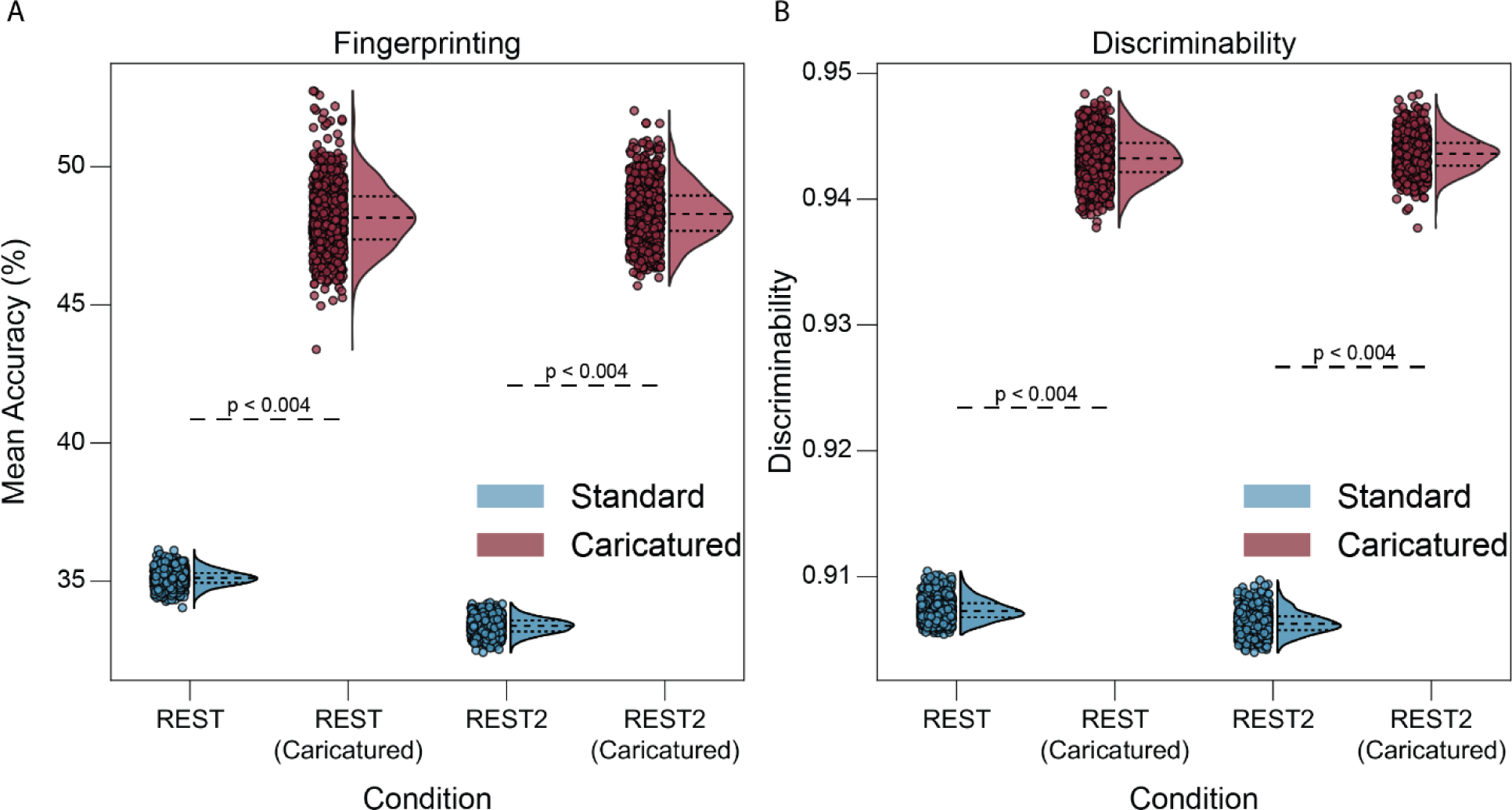
Multivariate reliability. Fingerprinting **(A)** and discriminability analysis **(B)** were performed for 1000 subsamples of individuals using pairs of LR and RL phase-encoded scans for each condition. Points represent mean fingerprinting accuracy **(A)** and discriminability **(B)** for each iteration, with the half violin plots demonstrating the distribution with the 1st, 2nd, and 3rd quartiles shown. P-values are shown for all relevant comparisons; they are denoted as “less than” a particular value when the resolution of the test could not be discerned below that value.

### 2.4 Univariate reliability decreases with caricaturing

Given the changes of within- and between-individual similarity and multivariate reliability, we investigated univariate (i.e., edge-level) reliability to understand how individual edges are changed after projection. Surprisingly, ICC was lower in Caricatured connectomes than in Standard ones. In the HCP dataset, Caricatured_HCP_ and Caricatured_CNP_ connectomes had a 25% decrease in ICC compared to Standard connectomes (p’s<0.0001; Bonferroni corrected; Figure 4A-B). In the TRT dataset (Figure 4C), ICC was 12% lower for the Caricatured_HCP_ connectomes and 9% lower for the Caricatured_CNP_ connectomes (p’s<0.0001; Bonferroni corrected).

**Figure 4:**
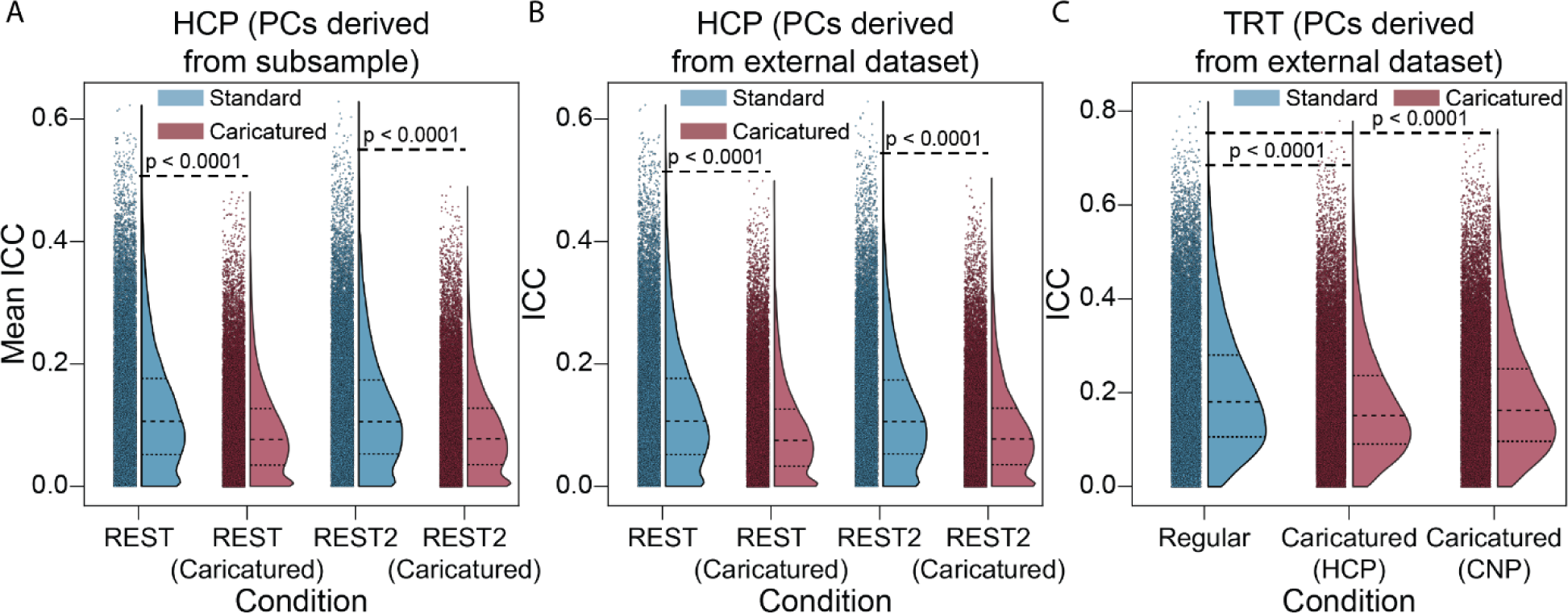
Intraclass correlation. **(A)** ICC was calculated for 1000 subsamples of participants using pairs of LR and RL phase encoded scans for each condition as the multiple measures per participant. Points represent the mean edge ICC across iterations with the half violin plots demonstrating the distribution with the 1st, 2nd, and 3rd quartiles shown. **(B)** ICC was calculated for LR and RL phase-encoded resting-state scans. Caricatured scans were projected onto PCs derived from the CNP dataset. **(C)** ICC was calculated in the TRT dataset using the run and session structure to partition the multiple measurements for each participant. Caricatured scans were projected onto PCs derived from the HCP or CNP datasets. P-values are shown for all relevant comparisons. We chose 0.0001 as the lowest bound above which to report p-values.

### 2.5 Caricatured connectomes improve resting-state predictive accuracy

Finally, as multivariate reliability provides an upper limit for prediction^30^, we examined whether improved CPM results for Caricatured connectomes accompanied the increased multivariate reliability. CPM models built from the Caricatured_HCP_ connectomes were significantly better than those from Standard connectomes (p’s<0.0001; Bonferroni corrected), explaining 40% more variance for age and 34% more variance for IQ (Figure 5A-B; **SI Figure 5; left panels**). Results were similar for Caricatured_CNP_ connectomes (Figure 5A-B; **SI Figure 5**; **right panels**), with a 21% average increase in explained variance for age and IQ (p’s≤0.0022; Bonferroni corrected). Sex classification was more accurate for Caricatured_HCP_ and Caricatured_CNP_ connectomes (p’s≤0.2808; Bonferroni corrected; Figure 5C). We also examined the multicollinearity of each model’s features to assess model interpretability. Across all phenotypes, multicollinearity was lower for models built on Caricatured_HCP_ connectomes (**SI Figures 6 and 7**). Additionally, models built on Caricatured_HCP_ connectomes had a similar or fewer number of features than those built on Standard connectomes (**SI Figure 8**). Thus, Caricatured connectomes increased CPM performance over Standard connectomes, partially by improving a connectome’s feature space and multivariate reliability.

**Figure 5:**
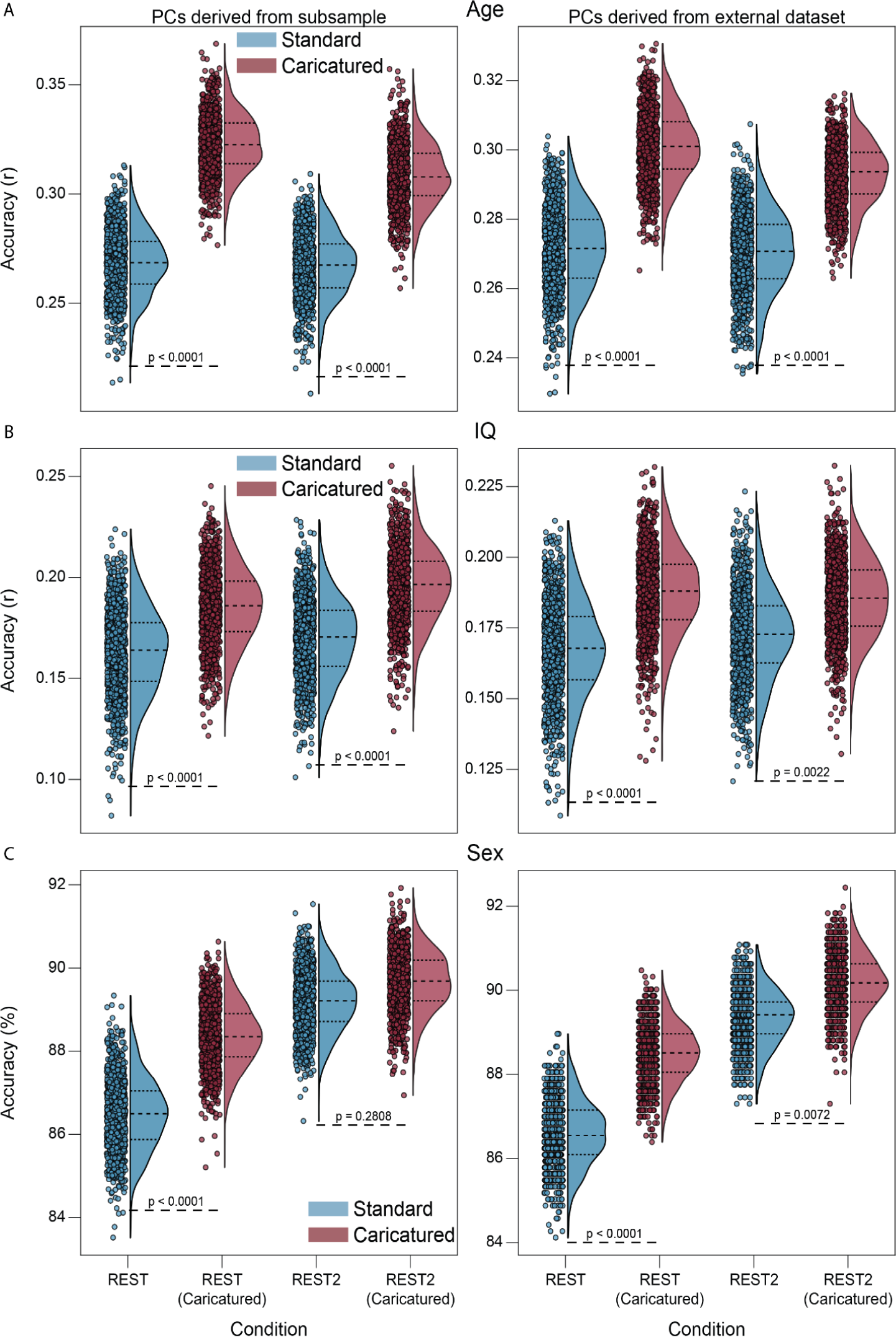
CPM prediction accuracy. Models were built for age **(A)**, IQ **(B)**, and sex **(C)** in the HCP. Caricatured connectomes were constructed using the HCP task data for the left and CNP task data for the right panels. For age and IQ, models were assessed via Pearson’s correlation between predicted and actual phenotype. For sex, models were evaluated via the percentage of participants correctly classified. In all plots, dots represent model performance for one of the 1000 randomizations of 10-fold cross-validation. P-values are shown for all relevant comparisons. We chose 0.0001 as the lowest bound above which to report p-values.

Overall, projecting resting-state data away from a manifold of task co-activation patterns improved prediction and multivariate reliability similar to task-based connectomes but mechanistically operated differently. Whereas task-based connectomes accomplish this improvement by increasing within- and between-individual similarity and edge-level reliability, caricaturing decreases these properties. Together, these results suggest that while moving resting-state data toward and away from tasks improves prediction and reliability, their underlying mechanisms are distinct.

## 3. Discussion

In this work, we introduced caricaturing—a method to project resting-state data to a subspace orthogonal to a manifold of co-activation patterns estimated from the task fMRI data. This projection removes linear combinations of these co-activation patterns from the resting-state data. It also formalizes the thought experiment from Finn et al., 2017^14^ and “caricatures” resting-state connectomes. These connectomes exhibited decreased between- and within-individual similarity and increased multivariate reliability and predictive utility. As caricaturing removes group-relevant task variance, it is an initial attempt to remove task-like co-activations from rest. Our results suggest that there is a useful signal beyond the dominating co-activations that drive resting-state functional connectivity. Therefore, caricaturing enhances individual differences in resting-state data and may better characterize the brain’s intrinsic functional architecture.

### 3.1 Co-activation events may hide useful signal

Resting-state functional connectivity can be difficult to interpret^31^. Early misconceptions about its nature painted a portrait of an intrinsic brain activity “baseline”^32^. Although refuted with evidence that there is a temporally unconstrained element of the signal associated with willful thought^33,34^, this only made the portrait more complex. If resting-state is some combination of task-like co-activation events^3,5–10^ and intrinsic functional architecture, then in theory, it should be possible to separate the two. Since our method projects resting-state data to a subspace orthogonal to a task-relevant manifold, the remaining signal may represent an intrinsic functional signal. Our results suggest that information about individual differences is present in this remaining signal, potentially at a greater degree than that found in the co-activation events. Nevertheless, these results warrant further study to determine what this signal represents.

### 3.2 Caricatured connectomes improve resting-state data

We used two datasets with different tasks to create the task manifold. Caricatured connectomes from both manifolds showed similar improvements in multivariate reliability and prediction. Thus, caricaturing appears general to the tasks and datasets used to create the manifold. As such, it is possible to use a large-scale, task-rich dataset to estimate the manifold and then apply it to any new or existing resting-state data. Thus, while task connectivity, naturalistic viewing paradigms, and “spotlighting” more generally offer exciting paths forward to study individual differences^35^, one need not throw out the baby with the bathwater. Existing resting-state data is widely available, and more will be collected. As such, approaches like ours that improve the utility of resting-state data are essential.

Relatedly, there may be other scenarios in which it is best to maximize within-individual similarity while minimizing between-individual similarity. So far, a single method has yet to accomplish this. Compared to resting-state connectomes, task-based connectomes increase both metrics^14^. By contrast, our method decreases both. Therefore, future work should determine whether novel approaches can be created to achieve the disentanglement of the two.

### 3.3 Caricaturing is similar to data prewhitening

Spotlighting and caricaturing are analogous to precoloring and prewhitening—two approaches from time series analysis, including in neuroimaging, to account for autocorrelations^36^. In precoloring, a large correlation is added to the time series to swamp out the unknown, existing autocorrelation. The total autocorrelation can be estimated as only the added component and input into the model to improve statistical inference. In spotlighting, tasks add a structure on top of rest, masking its unconstrained nature and improving downstream analyses (i.e., identification and prediction) in a conceptually similar way. However, precoloring also acts as a low-pass filter, which may remove signals of interest and degrade power^37^, similar to how tasks may obscure useful underlying signals. In contrast to precoloring, prewhitening estimates the true autocorrelation to remove it directly. Once removed, classic statistical inference is valid. Similarly, our caricaturing method estimates group-level, task-like co-activations and removes them from resting-state data to improve identification and prediction. In most applications, prewhitening is preferred to precoloring. However, prewhitening requires an accurate autocorrelation model, which can be difficult to determine with real data, similar to how it is difficult to estimate true task-like co-activation patterns. As with prewhitening and precoloring, future research will clarify the strengths and weaknesses of spotlighting and caricaturing and how they complement each other.

### 3.4 Multivariate reliability is separable from univariate reliability

Caricatured connectomes have greater multivariate but lower univariate reliability. While often going hand in hand, increasing multivariate does not guarantee an increase in univariate reliability^38^. They are distinct. Similarly, improving univariate reliability may not improve predictive utility^13,39,40^. Our results further explain this observation. Prediction methods are inherently multivariate, reflecting a pattern distributed across many features. Improving each feature’s reliability (i.e., univariate) may not make the overall pattern more reliable or, in turn, predictive. Therefore, methods for increasing multivariate reliability instead of univariate reliability are better equipped to improve predictive models.

### 3.5 Limitations

There are several limitations of our work. Although we project resting-state data away from a manifold of task co-activation patterns, it is unclear if task-relevant information is removed from the resting-state data. However, a couple of alternative explanations can be ruled out. First, the top PCs are unlikely to represent physiological noise. The PCs are defined at the group level, and heart rate, respiratory rate, and blood pressure would be averaged out across individuals. Second, the top PCs are unlikely to represent site- or scanner-related effects.

Results were generalizable across the datasets collected at different sites and with different scanners. Further, in line with previous research^25^, the PCs temporally track with task designs. Thus a portion of information in the PCs is certainly attributable to the task. Future research is needed to determine how much task-relevant information is present in resting-state and how it can be removed.

Additionally, we only used PCA to define the manifold. However, many nonlinear methods exist^7,41,42^ which may better identify task co-activation patterns. Further advances through nonlinear techniques may reveal complementary results.

### 3.6 Conclusion

In conclusion, we introduce a caricaturing method that projects resting-state fMRI data away from a manifold of task co-activation patterns, improving resting-state connectomes’ reliability and predictive utility. This method putatively removes task co-activation patterns. Our work suggests that the signal remaining after projection has increased reliability and utility over the standard resting-state signal. If resting-state combines intrinsic functional architecture and task-like co-activations, this signal may better represent this intrinsic functional architecture and is a topic for further study. Meanwhile, caricaturing can be applied to existing and future resting-state data to improve results in standard connectivity analyses.

## 4. Methods

### 4.1 Datasets

Three datasets were used in this work: the Human Connectome Project (HCP)^11^, the UCLA Consortium for Neuropsychiatric Phenomics (CNP)^12^, and the Yale test-retest dataset (TRT), which is composed of the publicly available TRTI^13^ and the private TRTII. HCP and CNP were chosen for their wide array of task-based data available in addition to resting-state data.

The TRT dataset was selected because participants were scanned four times across different days, yielding ideal data for assessing ICC and discriminability.

### 4.2 Processing

For the HCP data, we only used participants with data for each of the seven tasks (EMOTION, GAMBLING, LANGUAGE, MOTOR, RELATIONAL, SOCIAL, WM) and both resting-state scans (REST, REST2) for both the left-to-right (LR) and right-to-left (RL) phase encodings from the S1200 release. We also removed participants from analysis if the mean motion across all of their scans was greater than 0.1mm, if any scan’s mean motion was greater than 0.15mm, or if they were missing any data. Based on these criteria, 661 participants (males: 316, females: 345) remained. For the CNP data, we only used participants with data for each of the six tasks (BART, PAMENC, PAMRET, SCAP, STOPSIGNAL, TASKSWITCH). We further excluded participants if their mean motion across these scans was greater than 0.1mm, if the mean motion in any scan was greater than 0.15mm, or if any of their scans were missing data. From these criteria, we remained with 136 participants (males: 78, females: 58). For the TRT dataset, participants were scanned six times on four days. Since some participants were missing the sixth run for some days, we excluded the sixth run for all days and participants. Also, each scan was collected at rest for six minutes, but for some runs, scans were shorter and were still included in our analysis. The resulting data comprised 20 participants (males: 9, females: 11).

Consistent preprocessing steps were applied to all datasets. We started with the minimally preprocessed data for the HCP dataset^43^. For the CNP and TRTII data, skull-stripping was performed with OptiBet^44^. The data were then registered into common space. Motion correction was done with SPM8. The TRTI dataset was first skull-stripped in FSL^45^ and then registered into common space. Motion correction was performed in SPM5. The data were also iteratively smoothed to a 2.5mm Gaussian kernel equivalent^46,47^. For all datasets, further preprocessing was performed using BioImage Suite^48^. These steps included regressing 24 motion parameters, regressing the mean white matter, gray matter, and CSF time series, removing linear and quadratic trends, and applying a low-pass Gaussian filter (cutoff frequency ∼0.12 Hz for HCP, CNP, and TRTII and ∼0.19Hz for TRTI). For more detailed accounts of the CNP and TRTI datasets, see Gao et al., 2021^7^ and Noble et al., 2017^13^, respectively.

### 4.3 Caricaturing

In Caricaturing, we project resting-state data away from a task manifold. This method has two parts. The first is to define a task manifold from group-level, task fMRI. First, we temporally concatenate all task scans for individuals. Then, we perform principal component analysis (PCA). Each principal component (PC) is a common spatial activity pattern across tasks. The second part is to project resting-state data away from this manifold. First, we create a matrix of PCs excluding the top PCs (e.g., the first five, as in this work). This matrix is multiplied by its transpose to obtain the projection matrix. Next, we multiply the projection matrix and each time point from a resting-state scan, orthogonalizing them to the task manifold. Caricatured connectomes are created by correlating these orthogonalized time series.

#### Principal Component Estimation

Using fMRI time series data from one participant, the data can be represented as a matrix with dimensions 𝑡 × 𝑛, where 𝑡 is the number of frames in the scan and 𝑛 is the number of nodes in the atlas. To estimate the PCs, the time series for each node are z-scored, and the z-scored matrix is inputted into a PCA algorithm, ensuring that the algorithm considers the frames to be observations of the nodes, which are the variables.

The output is an 𝑛 × 𝑛 matrix 𝐿 of loadings, or PCs, which are orthonormal patterns of co-activation in the brain ordered by decreasing amount of variance explained in the data. To extend this framework to multiple scans, first, each time series is z-scored individually, and then they are concatenated along the time dimension. Thus, if using 𝑚 scans where each scan 𝑖 has 𝑡_𝑖_ frames, the final matrix to be input into the PCA algorithm will have dimensions (∑^𝑚^ 𝑡_𝑖_) × 𝑛. The resulting PCs are co-activation patterns that explain variance in the concatenated data.

#### Projection onto Principal Components

Using a time series matrix 𝑀 with dimensions 𝑡 × 𝑛 and a loading matrix 𝐿 with dimensions 𝑛 × 𝑛, we first choose a subset of PCs onto which we will project the time series. Let 𝑙 be vector of a subset of the integers from 1 to 𝑛 indicating which PCs will be used for projection. Then, we can create a new matrix 𝐿̂ where the 𝑖^𝑡ℎ^ column is equal to the (𝑙_𝑖_)^𝑡ℎ^ column of 𝐿 where 𝑙_𝑖_ is the 𝑖^𝑡ℎ^ element of 𝑙. We then create a projection matrix 𝑃𝑃 = 𝐿̂ × 𝐿̂𝑇, where 𝑇 indicates the transpose of a matrix. To project 𝑀 onto the desired PCs, we simply multiply it by the projection matrix to obtain 𝑀̂ = 𝑀 × 𝑃. The resulting matrix 𝑀̂ still retains the same dimensions as 𝑀 but now with only information from co-activation patterns that can be constructed by the desired PCs. Thus, the projected time series data can still be used for downstream connectomics analysis with the same dimensionality.

#### Implementation

The first five PCs obtained by concatenating time series across participants and tasks strongly reflected various aspects of the task structure^25^. Based on this result, we implemented our framework by projecting each participant’s z-scored resting-state time series onto the last 263 PCs of the task time series across multiple participants. Thus, we remove information from the top 5 group-level task PCs to remove dominating signals in the task data from the resting-state data.

### 4.4 Connectome Construction

Time series data were parcellated according to the Shen268 (268 nodes) atlas^49^, whereby the mean time course for each node was computed as the average of all voxel-level time series in that node. Connectomes were then constructed by taking the Fisher transform of the Pearson correlation between all pairs of node-wise time series. As a subsampling procedure was used in some analyses to ensure there was no data leakage between the data used to construct the PCs and the connectomes from which those PCs were projected away, we constructed Caricatured resting-state connectomes in the REST and REST2 HCP data for each subsample separately.

### 4.5 Downstream Metrics

We evaluated how caricaturing affects downstream connectome metrics. We assessed the connectomes constructed from projected resting-state time series (referred to as Caricatured connectomes) via within- and between-subject similarity, fingerprinting, discriminability, intraclass correlation (ICC), and connectome-based predictive modeling (CPM).

#### Similarity

To calculate similarity within and between individuals, we extracted and vectorized the upper triangle of the connectome. The within-individual similarities were computed as the correlation between vectorized connectome pairs of the same individual across scans. The between-individual similarities were calculated as the correlation between vectorized connectome pairs between different individuals.

#### Fingerprinting

We performed fingerprinting as described in Finn et al., 2015^50^. Given two groups of distinct connectomes that span the same participants, we labeled one group as the ‘Database’ and the other as the ‘Target Set’. For each connectome in the ‘Target Set’, the

Pearson correlation between that connectome and each in the ‘Database’ was calculated. The identity of the connectome in the ‘Database’ that corresponded to the highest correlation was assigned as the predicted identity of the current connectome in the ‘Target Set’. After repeating this for all connectomes in the ‘Target Set’, the fingerprinting accuracy for this label of ‘Database’ and ‘Target Set’ was calculated as the number of participants correctly identified divided by the number of participants. Perfect separability analysis–described in Noble et al., 2017^13^–is a simple extension to datasets with more than two scans per participant. From this, we can calculate the perfect separability rate (PSR), the percentage of scans for which all within-individual similarities are higher than any between-individual comparison.

#### Discriminability

Although fingerprinting serves as an excellent metric for participant identifiability, because it is binary in its methodology (i.e., correct vs. incorrect identification), it potentially leaves out information. Discriminability seeks to overcome this limitation by centering the method around the ratio of between-individual measurement distances that exceed within-individual measurement distances^30,51^. Furthermore, like perfect separability analysis, discriminability allows for any number of measurements per participant. We started by constructing a distance matrix between all pairs of measurements, using the correlation distance, or 1 minus Pearson’s r. Discriminability was calculated as the proportion of instances within-subject measurements were closer in distance than between-subject measurements across all possible combinations.

#### Intraclass Correlation

Whereas fingerprinting and discriminability are measures of multivariate reliability, that is, how reliable are connectomes as a whole, intraclass correlation (ICC) is a measure of univariate reliability, or how reliable are the edges of a connectome^52^. ICC^53^ is the fraction of variance due to the participant divided by the variance due to error. In the HCP data, there were only two measures per subject in each condition, so the variance components were estimated with a 2-way ANOVA. Symbolically, using subscripts s (subject), r (run), and e (residual) to represent the factors, this can be represented as *ICC*(*x*) = 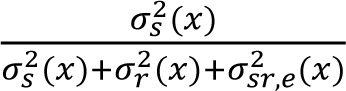, where 𝑥 is an edge in the connectome. In the TRT data, since the 20 scans per participant were partitioned by day and run, the variance components were estimated with a 3-way ANOVA. Here, adding d (day) to the factors, we get 𝐼(𝑥) = 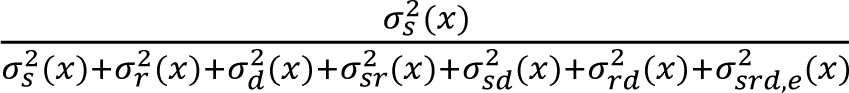. We set negative variance components (which were small in magnitude) to zero before computing ICC as in prior work^54^. For more information, see the shared code provided in the links below.

#### Connectome-Based Predictive Modeling

For our CPM analyses, we chose sex, fluid intelligence (IQ), and age to predict in the HCP dataset. These phenotypes are common in benchmarking analyses and typically have larger effect sizes. In all cases, models were built with 10-fold cross-validation where models were trained on 90% of the families and tested in 10% of the families in each fold. In each fold, feature selection was done to reduce the number of connectome edges used to build the model. Here, in the 90% of families used to build the model, edges were associated with the phenotype by either correlation (if continuous) or t-test (if binary). The resulting p-values for each edge were then observed and edges with a p-value less than 0.05 were used to build the model. For each phenotype, 1000 iterations of this 10-fold cross-validation were performed. For the continuous variables, the models were built using ridge regression. For sex, the models were built using linear support vector machine (SVM).

### 4.6 Statistics

This section provides specific details for all tests performed on the results of this research.

#### Similarity

For similarity analysis performed on the HCP data using the LR and RL phase encodings as the two scans per subject, tests assessing differences across the medians of within-subject similarity distributions were performed between REST and CaricaturedHCP REST, and REST2 and Caricatured_HCP_ REST2. We also performed tests to assess the same differences in medians of between-subject similarity distributions. The test performed is a paired, one-way non-parametric subtraction test performed in both directions whereby one distribution of median similarity is subtracted from the other, and 1 minus the proportion of differences that are greater than 0 is the resulting p-value. We then use Bonferroni correction to adjust the p-values across conditions (i.e., REST and REST2) and sub-analysis (i.e., within-subject and between-subject similarity). Thus, to be significant, a test must produce an uncorrected p-value less than 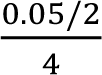. If an uncorrected p-value is returned as 0, we say it is less 4 than 0.001, since there are 1000 observations.

For the similarity analysis involving HCP connectomes projected onto the CNP PCs, we compared the full distributions of within-subject and between-subject similarity between REST and Caricatured_CNP_ REST, and REST2 and Caricatured_CNP_ REST2. A paired t-test was used here, and Bonferroni correction was applied across conditions and sub-analysis (multiplying the p-value by 4 to correct).

For the TRT similarity analysis, we compared the distribution of within-subject similarity in Standard resting-state connectomes to both Caricatured_HCP_ and Caricatured_CNP_ connectomes. The same was done for between-subject similarity. We again used the paired t-test with Bonferroni correction to account for the four tests performed.

#### Fingerprinting

For fingerprinting performed between LR and RL phase encodings for each scan condition, tests assessing differences in mean accuracy were performed between REST and Caricatured REST, and REST2 and Caricatured REST2. This was done via the same non-parametric subsampling test described above. Afterwards, Bonferroni correction was applied across these two tests, multiplying uncorrected p-values by 2 to correct. Thus, to be significant, a test must produce an uncorrected p-value less than 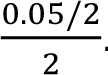. If an uncorrected p-value is 2 returned as 0, we say it is less than 0.001, since there are 1000 observations.

For the fingerprinting analysis involving HCP connectomes projected onto the CNP PCs, we compared accuracies between Caricatured_CNP_ connectomes and their standard counterparts. We performed a permutation test with 1000 permutations in which the labels for caricatured versus standard connectomes were shuffled with a probability of 0.5 for each scan to construct two distributions of null fingerprinting accuracies. The measured difference between REST and Caricatured_CNP_ REST accuracy and REST2 and Caricatured_CNP_ REST2 accuracy were compared to the distribution of differences between the constructed null accuracy distributions. Thus, the resulting uncorrected p-value in each case was 1 minus the proportion of times the empirical difference was greater than the null differences. If an uncorrected p-value was returned as zero, we stated that it was less than 0.001. Since this test was one-tailed and we compared two conditions, the resulting p-values underwent Bonferroni correction by multiplying them by four.

For the TRT perfect separability analysis, we compared the PSR in Standard resting-state connectomes to both Caricatured_HCP_ and Caricatured_CNP_ connectomes. For both comparisons, we used the permutation test described above. Again, the resulting p-values underwent Bonferroni correction by multiplying them by four. Importantly, statistical inference for PSR is not well-behaved, so the p-values are likely inaccurate. However, the discriminability analysis overcomes these limitations.

#### Discriminability

For the discriminability analysis using LR and RL phase encoded HCP connectomes, each subsample iteration yielded a single discriminability value. These were all pooled together to compare REST to Caricatured_HCP_ REST and REST2 to Caricatured_HCP_ REST2. The same non-parametric subsample test was used in each case, so to be significant, a test must produce an uncorrected p-value less than 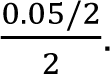. If an uncorrected p-value is 2 returned as 0, we say that it is less than 0.001, since there are 1000 observations.

To compare discriminability between REST and Caricatured_CNP_ REST and REST2 and Caricatured_CNP_ REST2 in the HCP dataset, we used the same type of permutation test as for fingerprinting accuracy. With 1000 iterations, if Caricatured discriminability was always greater, we defined the uncorrected p-value as less than 0.001. Using the Bonferroni method to correct for both tests and the fact that the test was one-tailed, we multiplied each p-value by 4.

Likewise, we used the same permutation test for the analysis where discriminability was computed in the TRT dataset. Comparisons were between Caricatured_HCP_ connectomes and the Standard TRT connectomes and between Caricatured_CNP_ connectomes and the Standard TRT connectomes. Bonferroni correction was applied by multiplying each p-value by 4.

#### Intraclass Correlation

For ICC calculated using the LR and RL phase encodings for each scan condition in the HCP dataset, 1000 subsample iterations were performed, yielding an ICC value for each edge in each iteration. The ICC values for each edge were averaged across iterations, yielding a mean ICC value for each edge in each scan condition. To compare REST to Caricatured_HCP_ REST and REST2 to Caricatured_HCP_ REST2, we used a Wilcoxon signed rank test and applied Bonferroni correction by multiplying the resulting p-values by 2.

For the ICC analysis in the HCP dataset, where caricatured connectomes used CNP-derived PCs, a single calculation yielded an ICC value for each edge in the connectome. Comparing REST to Caricatured_CNP_ REST and REST2 to Caricatured_CNP_ REST2, we used a Wilcoxon signed rank test and multiplied the resulting p-values by 2 to correct for multiple comparisons.

For the ICC analysis in the TRT dataset, two comparisons were performed using a Wilcoxon signed rank test. Edge ICC in the Caricatured_HCP_ connectomes and edge ICC in the Caricatured_CNP_ connectomes were compared to edge ICC in the Standard TRT connectomes. Bonferroni correction was applied by multiplying the resulting p-values by 2.

#### Connectome-Based Predictive Modeling

The correlation between actual and predicted phenotype assessed model performance for continuous variables. For both projections (i.e., Caricatured_HCP_ and Caricatured_CNP_), Caricatured REST was compared to Standard REST and Caricatured REST2 was compared to Standard REST2, using the 1000 subsample iterations to estimate the true model accuracy. Here, we used a corrected paired t-test similar to Nadeau and Bengio, 2003^55^ and referred to as the “corrected repeated k-fold CV test” in Bouckaert and Frank, 2004^56^. For each random subsample 𝑖, we calculated prediction accuracy for the caricatured data and the standard data, say 𝑎_𝑖_ and 𝑏_𝑖_. Letting the mean be 𝑚 = 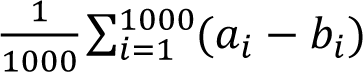 and the estimated variance be 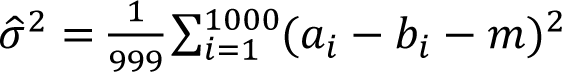, we arrive at the t-statistic 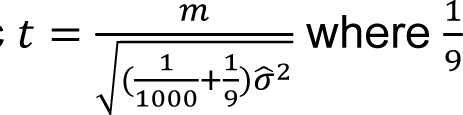 is an added correction factor to account for the lack of independence in sample pairs. This is input into the t-distribution with 999 degrees of freedom to compute the p-value. As this is a one-sided test to determine whether accuracy is greater for the caricatured connectomes, the p-value is multiplied by 2. Finally, Bonferroni correction is applied within each sub-analysis (i.e., age and IQ) and within each method for constructing PCs, so every p-value is again multiplied by 2 to correct for multiple comparisons. For the binary phenotype, model accuracy was assessed as the percentage of subjects correctly classified.

The same test was applied to compare REST to Caricatured REST and REST2 to Caricatured REST2, and the same statements regarding p-values and correction for multiple comparisons apply.

### 4.7 Data availability

The HCP data are publicly available on the ConnectomeDB database (https://db.humanconnectome.org/app/template/Login.vm). The UCLA CNP data can be obtained from the OpenfMRI database (https://openfmri.org/dataset/ds000030/). The TRTI data can be accessed publicly (https://fcon_1000.projects.nitrc.org/indi/retro/yale_trt.html). The TRTII data is currently unavailable for public access. Data used to generate the atlas parcellation can be accessed at http://fcon_1000.projects.nitrc.org/indi/retro/yale_hires.html.

### 4.8 Code availability

Code used for caricaturing can be found at https://github.com/RXRodriguez98/Caricature. Code used for fingerprinting and perfect separability analysis is available at https://github.com/SNeuroble/fingerprinting. Code used for discriminability analysis is available at https://github.com/RXRodriguez98/DiscriminabilityMATLAB. Code used for ICC calculation is available at https://github.com/SNeuroble/Multifactor_ICC. Code used for CPM can be found at https://github.com/YaleMRRC/CPM/tree/master and https://github.com/mattrosenblatt7/trust_connectomes/tree/main/utils/cpm_kfold_classification.

### 4.9 Ethics statement

All human subject data were collected previously (with informed consent) under the guidance of the local IRBs of the data collection sites. Yale Human Research Protection Program approved secondary analyses of these datasets.

## Supporting information

All Supplemental Information

## Acknowledgements

This work was supported by the National Institute of Mental Health [grant number R01MH121095] (obtained by D.S. and R. Todd Constable). R.X.R. was supported by the National Institute of General Medical Sciences [grant number 5T32GM100884-10]. S.N. was supported by the National Institute of Mental Health [grant number R00MH130894]. C.C.C. was supported by the Gruber Science Fellowship and the National Science Foundation Graduate Research Fellowship [grant number DGE-2139841]. Data were provided in part by the Human Connectome Project, WU-Minn Consortium (principal investigators, D. Van Essen and K. Ugurbil; 1U54MH091657) funded by the 16 US National Institutes of Health (NIH) institutes and centers that support the NIH Blueprint for Neuroscience Research; and by the McDonnell Center for Systems Neuroscience at Washington University. Data were also provided in part by the Consortium for Neuropsychiatric Phenomics (NIH Roadmap for Medical Research grants UL1-DE019580, RL1MH083268, RL1MH083269, RL1DA024853, RL1MH083270, RL1LM009833, PL1MH083271, and PL1NS062410). CNP data was obtained from the OpenfMRI database. Its accession number is ds000030.

## Author Contributions

R.X.R. conceptualized the study with guidance from D.S. and S.N. R.X.R. performed the analysis with support from C.C.C., S.N., and D.S. S.N. designed, collected, and preprocessed the Yale Test-Retest dataset. D.S. and S.N. provided guidance on result interpretation. R.X.R. wrote the manuscript with contributions from D.S., S.N., and C.C.C., and comments from all authors.

## Materials & Correspondence

Correspondence to Raimundo X. Rodriguez or Dustin Scheinost.

## References

1. Allan, T. W. et al. Functional Connectivity in MRI Is Driven by Spontaneous BOLD Events. PLOS ONE 10, e0124577 (2015).

2. Gonzalez-Castillo, J., Kam, J. W. Y., Hoy, C. W. & Bandettini, P. A. How to Interpret Resting-State fMRI: Ask Your Participants. J. Neurosci. 41, 1130–1141 (2021).

3. Petridou, N., Gaudes, C. C., Dryden, I. L., Francis, S. T. & Gowland, P. A. Periods of rest in fMRI contain individual spontaneous events which are related to slowly fluctuating spontaneous activity. Human Brain Mapping 34, 1319–1329 (2013).

4. Tagliazucchi, E., Siniatchkin, M., Laufs, H. & Chialvo, D. R. The Voxel-Wise Functional Connectome Can Be Efficiently Derived from Co-activations in a Sparse Spatio-Temporal Point-Process. Frontiers in Neuroscience 10, (2016).

5. Zamani Esfahlani, F., et al. High-amplitude cofluctuations in cortical activity drive functional connectivity. Proceedings of the National Academy of Sciences 117, 28393–28401 (2020).

6. Chen, R. H., Ito, T., Kulkarni, K. R. & Cole, M. W. The Human Brain Traverses a Common Activation-Pattern State Space Across Task and Rest. Brain Connectivity 8, 429–443 (2018).

7. Gao, S., Mishne, G. & Scheinost, D. Nonlinear manifold learning in functional magnetic resonance imaging uncovers a low-dimensional space of brain dynamics. Human Brain Mapping 42, 4510–4524 (2021).

8. Gonzalez-Castillo, J. et al. Imaging the spontaneous flow of thought: Distinct periods of cognition contribute to dynamic functional connectivity during rest. NeuroImage 202, 116129 (2019).

9. Kim, D., Livne, T., Metcalf, N. V., Corbetta, M. & Shulman, G. L. Spontaneously emerging patterns in human visual cortex and their functional connectivity are linked to the patterns evoked by visual stimuli. Journal of Neurophysiology 124, 1343–1363 (2020).

10. Tusche, A., Smallwood, J., Bernhardt, B. C. & Singer, T. Classifying the wandering mind: Revealing the affective content of thoughts during task-free rest periods. NeuroImage 97, 107–116 (2014).

11. Van Essen, D. C. et al. The WU-Minn human connectome project: an overview. Neuroimage 80, 62–79 (2013).

12. Poldrack, R. A. et al. A phenome-wide examination of neural and cognitive function. Sci Data 3, 160110 (2016).

13. Noble, S. et al. Influences on the test–retest reliability of functional connectivity MRI and its relationship with behavioral utility. Cerebral cortex 27, 5415–5429 (2017).

14. Finn, E. S. et al. Can brain state be manipulated to emphasize individual differences in functional connectivity? NeuroImage 160, 140–151 (2017).

15. Noble, S., Scheinost, D. & Constable, R. T. A decade of test-retest reliability of functional connectivity: A systematic review and meta-analysis. NeuroImage 203, 116157 (2019).

16. Greene, A. S., Gao, S., Scheinost, D. & Constable, R. T. Task-induced brain state manipulation improves prediction of individual traits. Nat Commun 9, 2807 (2018).

17. Greene, A. S., Gao, S., Noble, S., Scheinost, D. & Constable, R. T. How Tasks Change Whole-Brain Functional Organization to Reveal Brain-Phenotype Relationships. Cell Reports 32, 108066 (2020).

18. Jiang, R. et al. Task-induced brain connectivity promotes the detection of individual differences in brain-behavior relationships. NeuroImage 207, 116370 (2020).

19. Zhao, W. et al. Task fMRI paradigms may capture more behaviorally relevant information than resting-state functional connectivity. NeuroImage 270, 119946 (2023).

20. Finn, E. S. & Bandettini, P. A. Movie-watching outperforms rest for functional connectivity-based prediction of behavior. NeuroImage 235, 117963 (2021).

21. Vanderwal, T. et al. Individual differences in functional connectivity during naturalistic viewing conditions. NeuroImage 157, 521–530 (2017).

22. Wang, J. et al. Test–retest reliability of functional connectivity networks during naturalistic fMRI paradigms. Human Brain Mapping 38, 2226–2241 (2017).

23. Yoo, K. et al. A cognitive state transformation model for task-general and task-specific subsystems of the brain connectome. NeuroImage 257, 119279 (2022).

24. Yoo, K. et al. A brain-based general measure of attention. Nat Hum Behav 6, 782–795 (2022).

25. Shine, J. M. et al. Human cognition involves the dynamic integration of neural activity and neuromodulatory systems. Nature neuroscience 22, 289–296 (2019).

26. Airan, R. D. et al. Factors affecting characterization and localization of interindividual differences in functional connectivity using MRI. Human Brain Mapping 37, 1986–1997 (2016).

27. Birn, R. M. et al. The effect of scan length on the reliability of resting-state fMRI connectivity estimates. NeuroImage 83, 550–558 (2013).

28. Laumann, T. O. et al. Functional system and areal organization of a highly sampled individual human brain. Neuron 87, 657–670 (2015).

29. Mueller, S. et al. Reliability correction for functional connectivity: Theory and implementation. Human Brain Mapping 36, 4664–4680 (2015).

30. Bridgeford, E. W. et al. Eliminating accidental deviations to minimize generalization error and maximize replicability: Applications in connectomics and genomics. PLoS computational biology 17, e1009279 (2021).

31. Biswal, B., Zerrin Yetkin, F., Haughton, V. M. & Hyde, J. S. Functional connectivity in the motor cortex of resting human brain using echo-planar MRI. Magnetic resonance in medicine 34, 537–541 (1995).

32. Stark, C. E. L. & Squire, L. R. When zero is not zero: The problem of ambiguous baseline conditions in fMRI. Proceedings of the National Academy of Sciences 98, 12760–12766 (2001).

33. Buckner, R. L., Krienen, F. M. & Yeo, B. T. T. Opportunities and limitations of intrinsic functional connectivity MRI. Nat Neurosci 16, 832–837 (2013).

34. Morcom, A. M. & Fletcher, P. C. Does the brain have a baseline? Why we should be resisting a rest. NeuroImage 37, 1073–1082 (2007).

35. Finn, E. S. Is it time to put rest to rest? Trends in Cognitive Sciences 25, 1021–1032 (2021).

36. Poldrack, R. A., Mumford, J. A. & Nichols, T. E. Handbook of Functional MRI Data Analysis. (Cambridge University Press, 2011).

37. Monti, M. Statistical Analysis of fMRI Time-Series: A Critical Review of the GLM Approach. Frontiers in Human Neuroscience 5, (2011).

38. Camp, C. C., Noble, S., Scheinost, D., Stringaris, A. & Nielson, D. M. Test-Retest Reliability of Functional Connectivity in Adolescents With Depression. Biological Psychiatry: Cognitive Neuroscience and Neuroimaging 9, 21–29 (2024).

39. Shirer, W. R., Jiang, H., Price, C. M., Ng, B. & Greicius, M. D. Optimization of rs-fMRI Pre-processing for Enhanced Signal-Noise Separation, Test-Retest Reliability, and Group Discrimination. NeuroImage 117, 67–79 (2015).

40. Strother, S. et al. Optimizing the fMRI data-processing pipeline using prediction and reproducibility performance metrics: I. A preliminary group analysis. NeuroImage 23, S196– S207 (2004).

41. Busch, E. L. et al. Multi-view manifold learning of human brain-state trajectories. Nat Comput Sci 3, 240–253 (2023).

42. Ye, J. et al. Altered Brain Dynamics Across Bipolar Disorder and Schizophrenia During Rest and Task Switching Revealed by Overlapping Brain States. Biological Psychiatry 94, 580– 590 (2023).

43. Glasser, M. F. et al. The minimal preprocessing pipelines for the Human Connectome Project. NeuroImage 80, 105–124 (2013).

44. Lutkenhoff, E. S. et al. Optimized brain extraction for pathological brains (optiBET). PloS one 9, e115551 (2014).

45. Smith, S. M. Fast robust automated brain extraction. Human Brain Mapping 17, 143–155 (2002).

46. Friedman, L., Glover, G. H., Krenz, D., Magnotta, V. & Birn, T. F. Reducing inter-scanner variability of activation in a multicenter fMRI study: role of smoothness equalization. Neuroimage 32, 1656–1668 (2006).

47. Scheinost, D., Papademetris, X. & Constable, R. T. The impact of image smoothness on intrinsic functional connectivity and head motion confounds. Neuroimage 95, 13–21 (2014).

48. Joshi, A. et al. Unified framework for development, deployment and robust testing of neuroimaging algorithms. Neuroinformatics 9, 69–84 (2011).

49. Shen, X., Tokoglu, F., Papademetris, X. & Constable, R. T. Groupwise whole-brain parcellation from resting-state fMRI data for network node identification. Neuroimage 82, 403–415 (2013).

50. Finn, E. S. et al. Functional connectome fingerprinting: identifying individuals using patterns of brain connectivity. Nat Neurosci 18, 1664–1671 (2015).

51. Wang, Z., Bridgeford, E., Wang, S., Vogelstein, J. T. & Caffo, B. Statistical analysis of data repeatability measures. arXiv preprint arXiv:2005.11911 (2020).

52. Noble, S., Scheinost, D. & Constable, R. T. A guide to the measurement and interpretation of fMRI test-retest reliability. Current opinion in behavioral sciences 40, 27 (2021).

53. Shrout, P. E. & Fleiss, J. L. Intraclass correlations: uses in assessing rater reliability. Psychological bulletin 86, 420 (1979).

54. Shavelson, R. J., Baxter, G. P. & Gao, X. Sampling variability of performance assessments. Journal of educational Measurement 30, 215–232 (1993).

55. Nadeau, C. & Bengio, Y. Inference for the Generalization Error. Machine Learning 52, 239– 281 (2003).

56. Bouckaert, R. R. & Frank, E. Evaluating the replicability of significance tests for comparing learning algorithms. in Pacific-Asia conference on knowledge discovery and data mining 3–12 (Springer, 2004).

