## Supplementary material for "Connectome caricatures: removing large-amplitude co-activation patterns in resting-state fMRI emphasizes individual differences": All Supplemental Information

*SI 1.1 Method replication and validation*

For each task in the HCP dataset, we compared PC time series to the task block regressors (**SI Figure 1A-B**), as was performed in Shine et al., 2019[1]. We created these subject-specific regressors by noting the start and stop times for each task block and convolving the block structure with the canonical hemodynamic response function defined in SPM12. Critically, the LANGUAGE task did not have any “rest” blocks, so we chose the auditory language task to serve as the rest time points and the math portion to serve as the task block, in line with Shine et al., 2019[1]. We constructed task block regressors for each task scan, split by phase encoding. We also concatenated all task block regressors for a given phase encoding in each subject to get the “All Tasks” regressor. Lastly, we took the absolute value of the derivative of the “All Tasks” regressors. One subject was missing task regressor information and was excluded from this analysis.

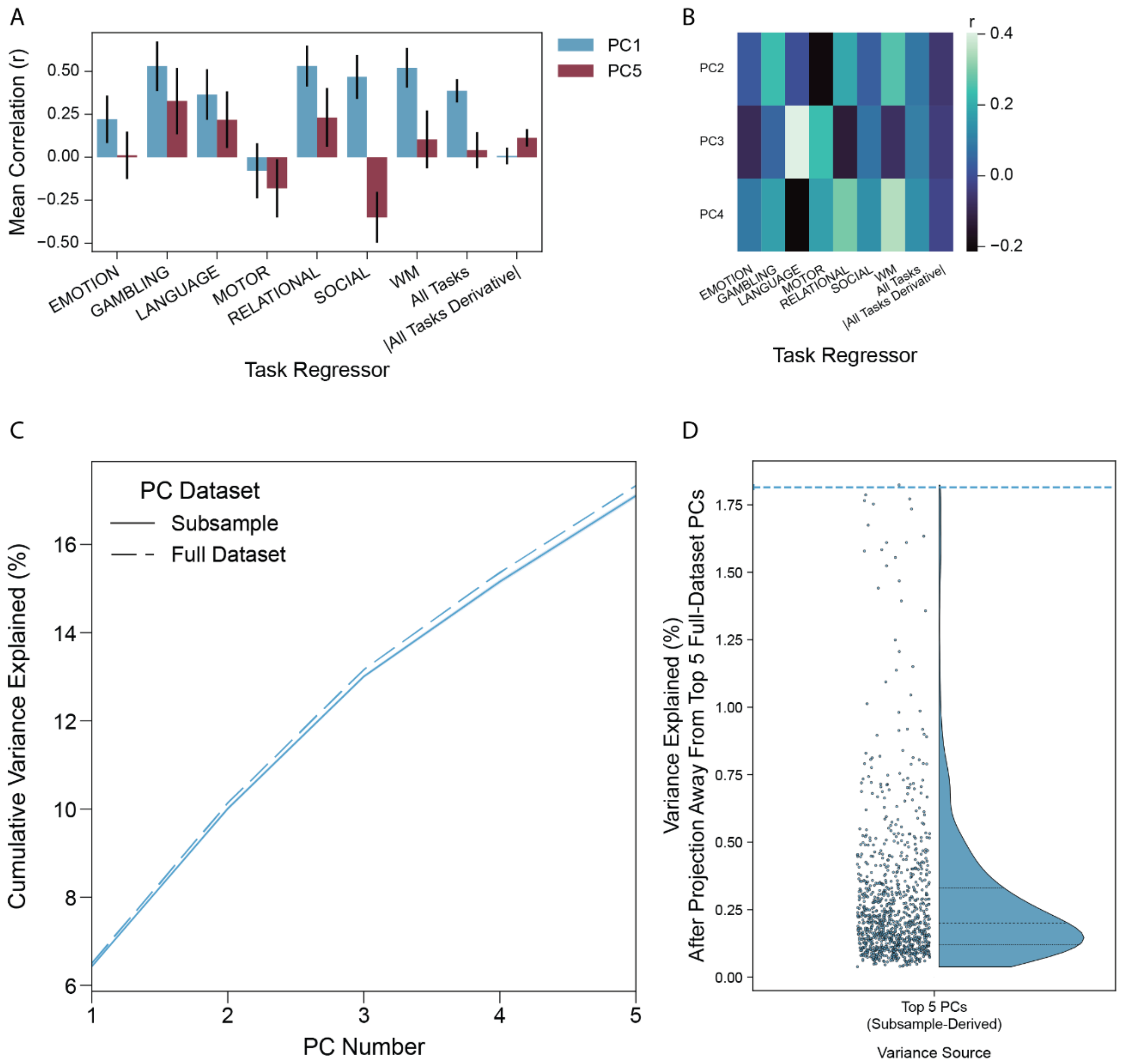


**SI Figure 1:** PCA replication and validation. **(A)** PCs were constructed by concatenating all task scans for every subject in the HCP dataset and performing PCA. Time series associated with the first and fifth PCs were correlated with task block regressors. Colors represent which PC time series was used in the correlation. The bars represent the mean correlation (across phase encodings and all subjects) for each correlation pair with error bars indicating one standard deviation. **(B)** Time series associated with the second, third, and fourth PCs were correlated with task block regressors. Colors represent the mean correlation across phase encodings and subjects. **(C)** PCs were constructed either in task scans for all HCP subjects (Full Dataset) or in 1000 subsamples of subjects from 10 families (Subsample). Their respective line plots indicate the cumulative variance explained by the first five PCs in the full dataset. The Subsample line also has an error band around it indicating the standard deviation of the measure across the 1000 iterations. **(D)** The full concatenated time series was projected away from the top five full-dataset PCs. Then, for each of the 1000 iterations, the amount of additional variance explained by the top five subsample-derived PCs was calculated. Points represent these values and the violin plot visualizes their distribution with the 1st, 2nd, and 3rd quartiles shown. The dashed line indicates the amount of variance explained by the sixth full-dataset PC.

When evaluating connectome quality in the same dataset in which we constructed the PCs, we used a subsampling procedure whereby subjects from 10 families (to control for family structure) were used to construct the PCs and the remaining subjects were used in the downstream analyses. This procedure was repeated 1000 times with different left-out families to ensure robust results without the risk of data leakage. To validate our subsampling procedure, we sought to verify that the top five subsample-derived PCs spatially and statistically recapitulated the top five full-dataset PCs. Here, we considered the full-dataset PCs as a ground truth and the subsample-derived PCs as estimates of the ground truth. For each iteration of the subsampling procedure, we calculated the cumulative percentage of variance explained by the top five subsample-derived PCs in the full concatenated dataset. Similarly, we calculated the cumulative percentage of variance explained by the top five full-dataset PCs in the same dataset. The cumulative percentage of variance explained by the subsample-derived PCs was similar to the percentage of variance explained by the full-dataset PCs (**SI Figure 1C**). This suggests that they behave in a statistically similar manner. Next, we considered whether the subsample-derived PCs and the full-dataset PCs explained variance from the same sources. We first projected the full concatenated dataset away from the top five full-dataset PCs. We then calculated the amount of variance explained in the projected time series by the top five subsample-derived PCs and compared it to the variance explained by the sixth full-dataset PC (**SI Figure 1D**). We observed that the variance explained by the sixth full-dataset PC was significantly higher than that explained by the top five subsample-derived PCs after the projection ($p=0.002$). Since, by construction, the sixth subsample-derived PC explains the next most amount of variance after the first five in the full dataset, this observation indicates that the top five subsample-derived PCs are a good spatial estimate of the ground truth.

The test used in **SI Figure 1D** was a paired, one-way non-parametric subtraction test. We tested in both directions. The distribution of variance explained was subtracted from the variance explained by the sixth full-dataset PC. One minus the proportion of differences that are greater than 0 is the resulting p-value. Thus, to be significant, a test must produce an uncorrected p-value less than 0.025.

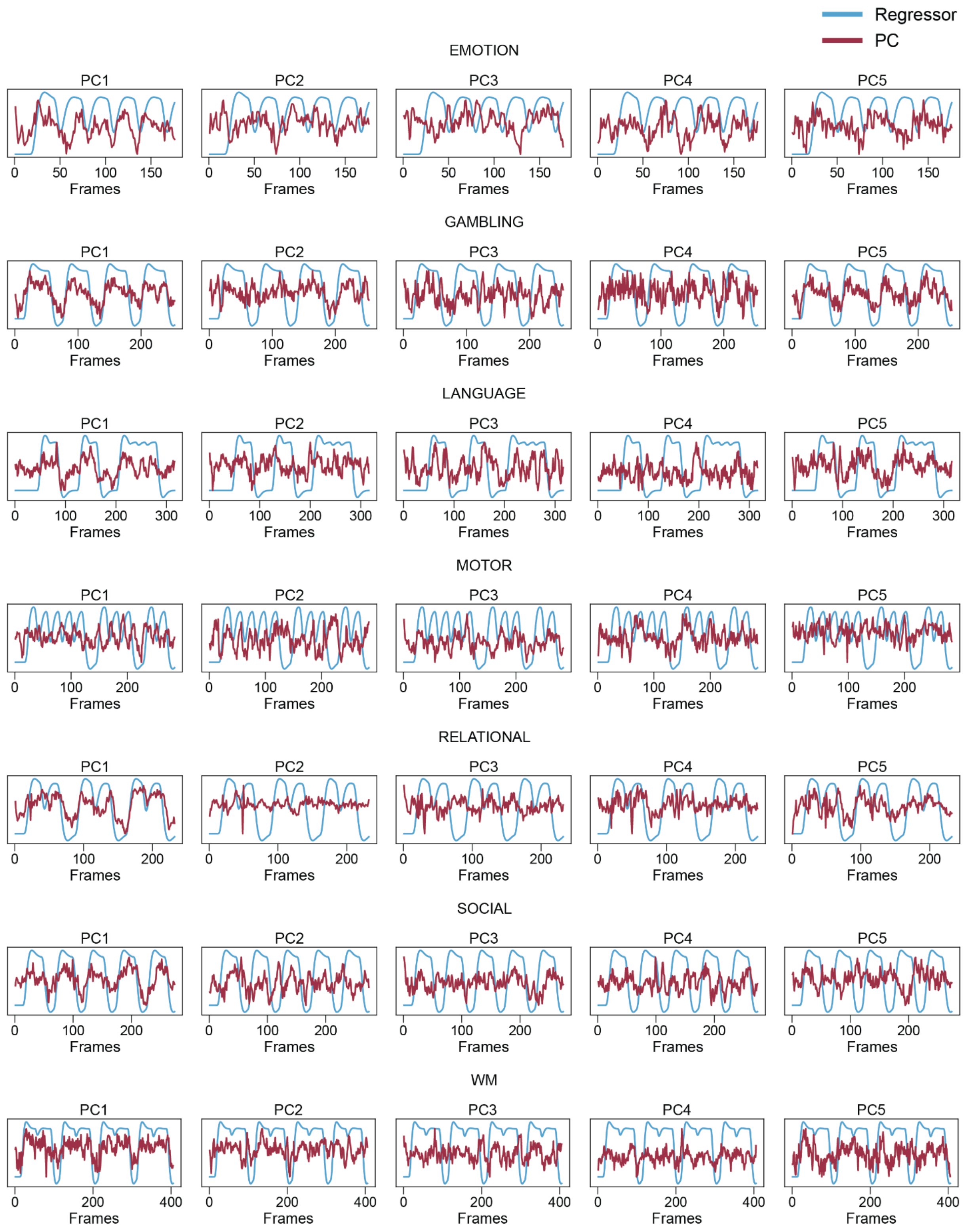


**SI Figure 2.** Task block regressor overlap with PC time series. Task block regressors were created by convolving the HRF with the block structure of each task, and these were compared to the PC time series. Results are shown for the LR phase encoding in the subject whose PC1 time series was most correlated with the full concatenated task block regressor. Here, this is broken down by task type and PC number to show how each PC tracks each task individually. All PC time series are normalized to range from 0 to 1 for visualization purposes.

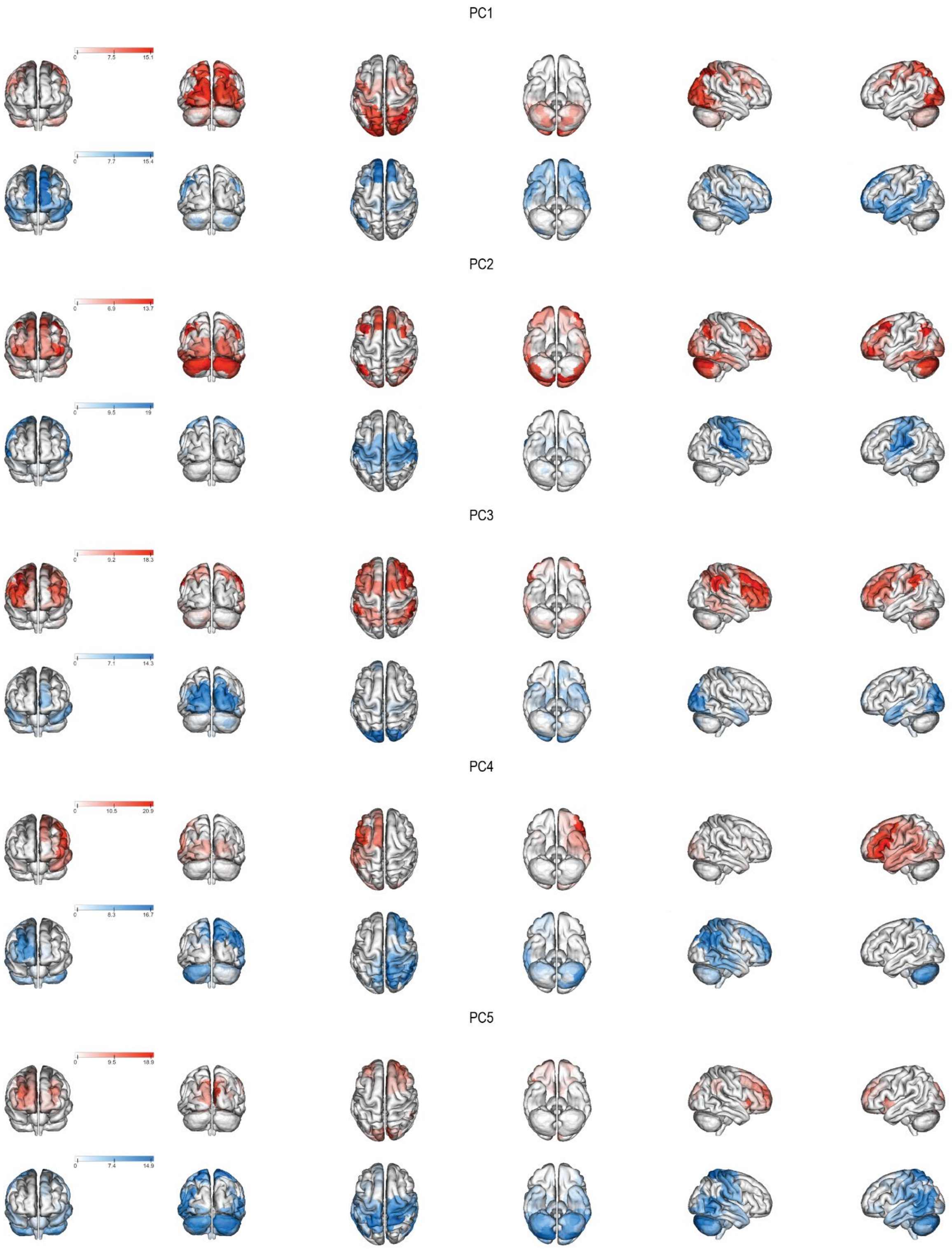


**SI Figure 3.** PC spatial distribution. Each figure shows the weight for every node for a given PC, where red indicates a positive weight and blue indicates a negative weight. PC weights were scaled by 100 to have a fuller color bar. Visualization was performed using BioImage Suite at https://bioimagesuiteweb.github.io/webapp/.

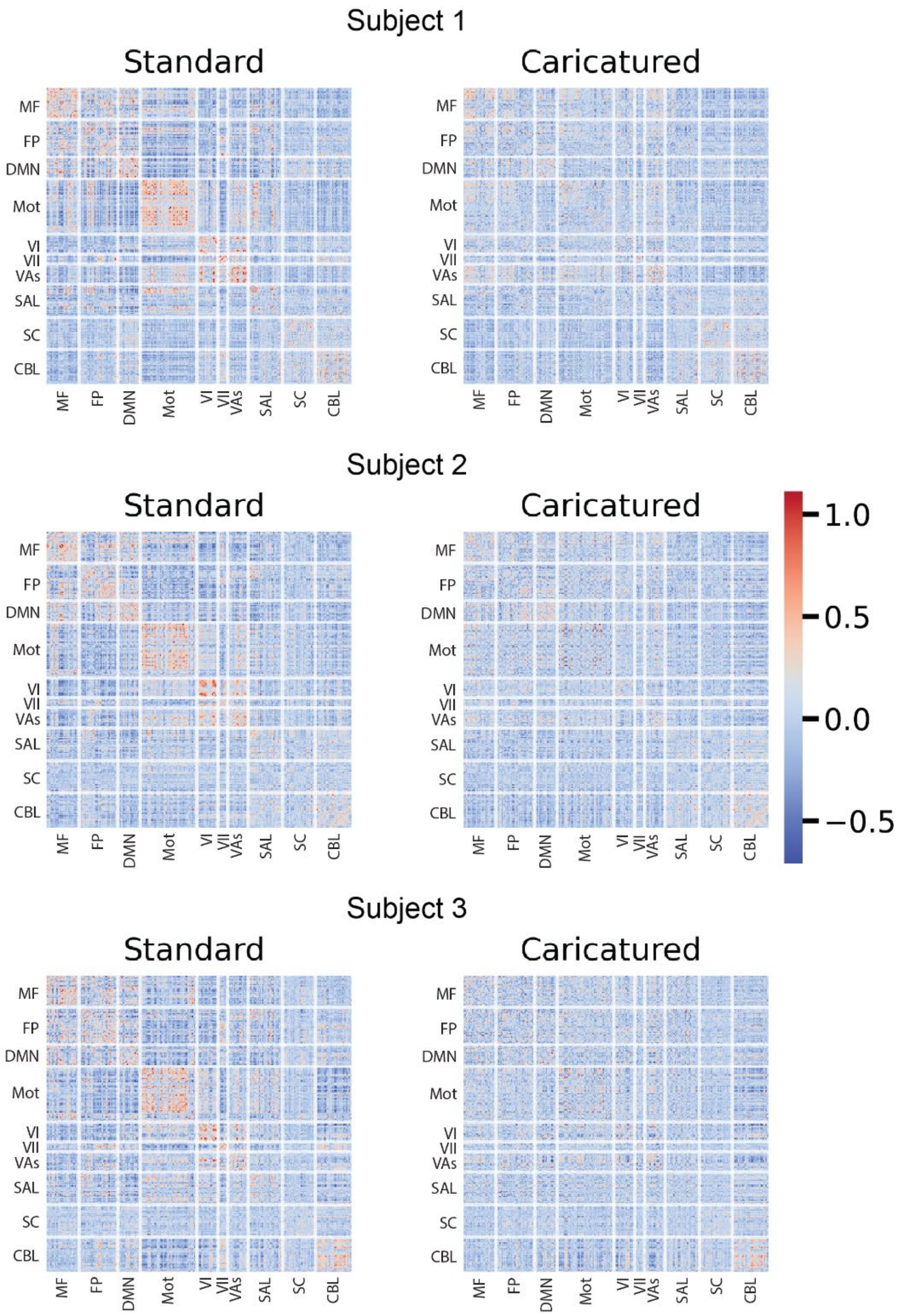


**SI Figure 4.** Example connectomes. To show how the PC projection procedure affects the resting-state connectomes, we show standard and caricatured connectomes for three subjects. All connectomes are reorganized and plotted by subnetwork[2,3].

*SI 1.2 Multivariate reliability improves using multiple datasets*

As some of the multivariate reliability analyses resulted in single numbers, we provide tables to display the results. Fingerprinting was performed in the HCP dataset with PCs constructed in the CNP dataset (**SI Table 1**). Using Caricatured$_{CNP}$ connectomes resulted in a 42% increase in accuracy compared to standard connectomes.

| Scan Condition | Connectome Type | Fingerprinting Accuracy (%) |
|---|---|---|
| REST | Standard | 34.95 |
| | $\text{Caricatured}_{CNP}$ | 48.18 |
| REST2 | Standard | 33.21 |
| | $\text{Caricatured}_{CNP}$ | 48.34 |

**SI Table 1.** HCP fingerprinting in Standard vs $\text{Caricatured}_{CNP}$ connectomes. Fingerprinting was performed using pairs of LR and RL phase-encoded scans for each condition. The resulting accuracy is the average across using each phase-encoding as the 'Database' and 'Target Set'.

Perfect separability rate (PSR) was calculated in the TRT dataset with PCs constructed for projection in both the HCP and CNP datasets (**SI Table 2**). Using $\text{Caricatured}_{HCP}$ connectomes resulted in a 633% increase in PSR, and using $\text{Caricatured}_{CNP}$ connectomes resulted in a 383% increase in PSR.

| Dataset | Connectome Type | PSR |
|---|---|---|
| TRT | Standard | 1.5% |
| | $\text{Caricatured}_{HCP}$ | 11% |
| | $\text{Caricatured}_{CNP}$ | 7.25% |

**SI Table 2.** PSR analysis in the TRT dataset. PSR was calculated using all 20 scans available per subject.

Likewise, discriminability analysis was performed in the HCP dataset with PCs constructed in the CNP dataset and in the TRT dataset with PCs constructed either in the HCP or CNP dataset (**SI Table 3**). In the HCP dataset, using $\text{Caricatured}_{CNP}$ connectomes resulted in a 4% increase in discriminability. In the TRT dataset, using Caricatured connectomes resulted in a 1% increase in discriminability.

| Dataset | Connectome Type | Discriminability |
|---|---|---|
| HCP REST | Standard | 0.9074 |
| | $Caricatured_{CNP}$ | 0.9414 |
| HCP REST2 | Standard | 0.9063 |
| | $Caricatured_{CNP}$ | 0.9453 |
| TRT | Standard | 0.9704 |
| | $Caricatured_{HCP}$ | 0.9842 |
| | $Caricatured_{CNP}$ | 0.9800 |

**SI Table 3.** Discriminability using an external dataset to construct PCs. Discriminability analysis was performed in the HCP dataset using LR and RL phase-encoded scans for each condition and in the TRT dataset using all 20 scans available per subject.

*SI 1.3 CPM mean squared error (MSE) accuracy and feature space quality*

To complement the CPM accuracy analyses using correlation for the continuous phenotypes, we also assessed model accuracy via MSE (**SI Figure 5**), where a lower MSE indicates a superior model. For age prediction in the HCP dataset using $Caricatured_{HCP}$ connectomes (**SI Figure 5A**; left panel), the MSE decreased by 2% (p’s<0.0001; Bonferroni corrected). For IQ prediction (**SI Figure 5B**; left panel), the MSE decreased by 1% (p’s<0.0001; Bonferroni corrected). We repeated both of these analyses instead using $Caricatured_{CNP}$ connectomes. For age prediction (**SI Figure 5A**; right panel), MSE decreased by 1% when using Caricatured connectomes. The difference between Standard and Caricatured performance was significant (p’s<0.0001; Bonferroni corrected). Likewise, for IQ prediction (**SI Figure 5B**; right panel), MSE decreased by 1%. This difference was significant (p’s<0.004; Bonferroni corrected). P-values were calculated using the same corrected paired t-test used to evaluate correlation-based accuracy. Corrections for multiple comparisons were applied analogously.

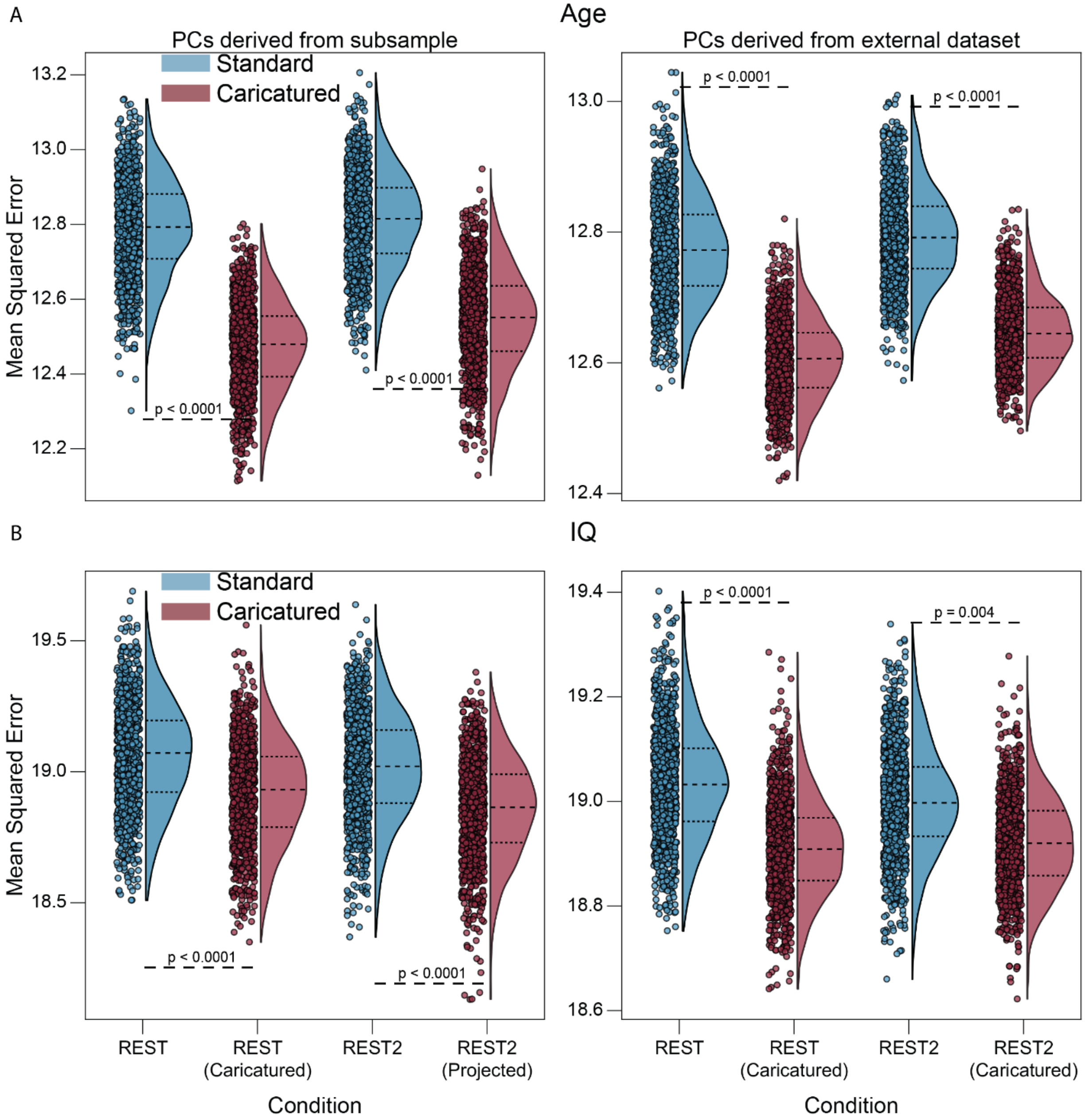


**SI Figure 5:** CPM mean squared error (MSE). Models were built for age **(A)** and IQ **(B)** in the HCP LR data in 1000 iterations. For the left panels, a subset of subjects were left out in each iteration to construct the PCs onto which resting-state scans of the remaining subjects were projected. For the right panels, the PCs were constructed using the CNP dataset. Models were assessed via MSE. The dots represent individual MSE for a given iteration, and the violin plots demonstrate the distribution of those points. P-values are shown for all relevant comparisons. We chose 0.0001 as the lowest bound above which to report p-values.

We also investigated the quality of the models built for each phenotype by observing properties of the feature space. Specifically, we assessed whether the selected features had substantial multicollinearity, which would indicate a less interpretable model (**SI Figures 6-7**), as well as how many features were selected (**SI Figure 8**). To assess multicollinearity, we used PCA on the feature space and calculated both the percentage of PCs needed to explain 80% of the feature space variance (**SI Figure 6**) as well as the normalized Shannon entropy using the percentage of variance explained by each PC as the probability distribution (**SI Figure 7**). In both cases, lower values would indicate less multicollinearity among the features.

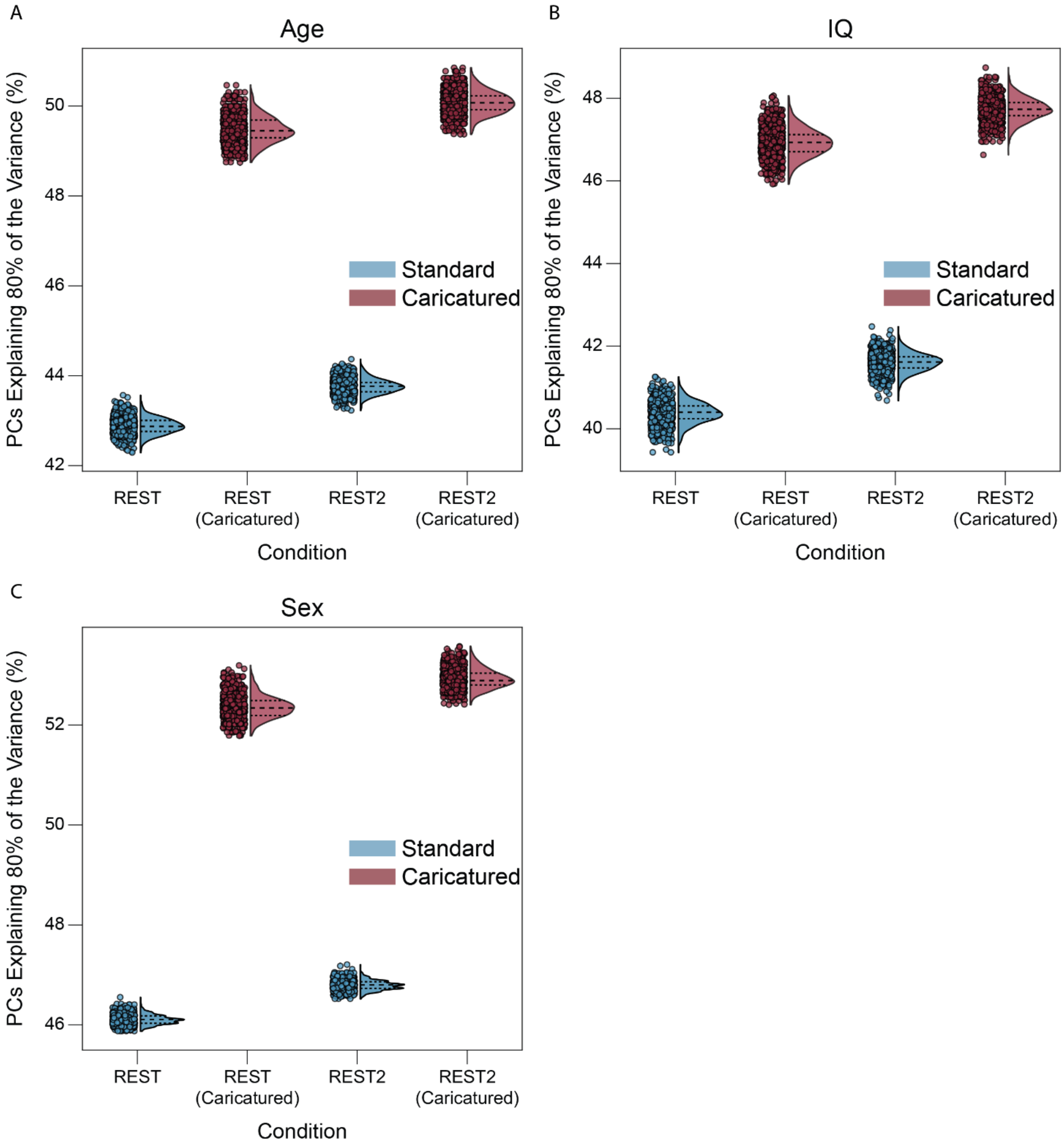


**SI Figure 6:** Feature multicollinearity measured by percentage of feature space PCs needed to explain 80% of the variance. To estimate the features used in each of the 1000 models built in the CPM process, we iterated through each of the subsamples and found the p-value for the association between the phenotype and each edge across subjects. As the p-value threshold for the models was 0.05, only edges with lower p-values were selected as features. Vectors of edge values across subjects were created for the features surviving the threshold. These were then z-scored and input into PCA such that the PCs were coefficients for linear combinations of features. Each point represents the percentage of PCs needed to explain 80% of the feature space for that iteration, and the violin plots show the distributions. This process was done for phenotypes age **(A)**, IQ **(B)**, and sex **(C)**.

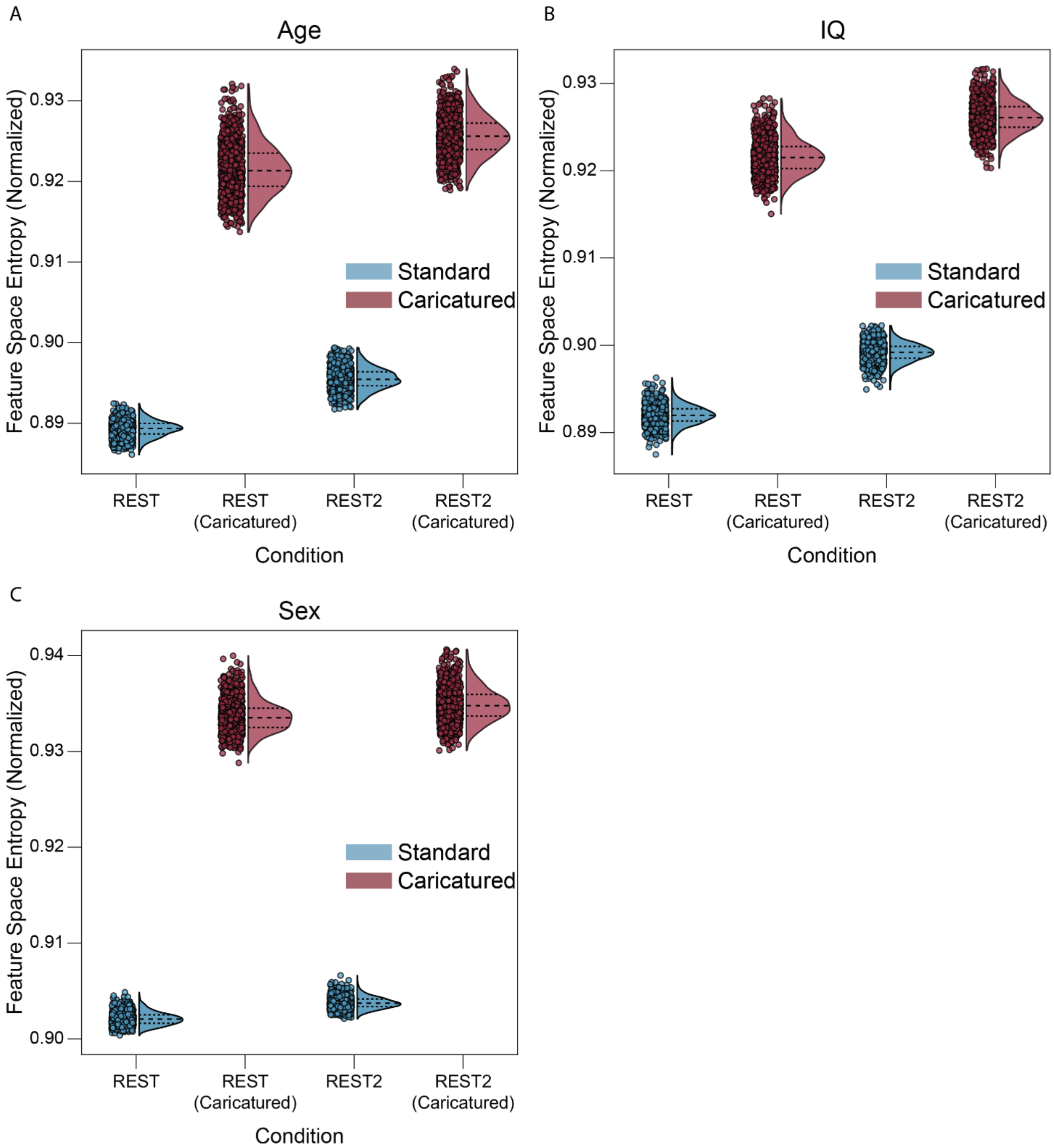


**SI Figure 7:** Feature multicollinearity measured by entropy of feature space PCs. To estimate the features used in each of the 1000 models built in the CPM process, we iterated through each of the subsamples and found the p-value for the association between the phenotype and each edge across subjects. As the p-value threshold for the models was 0.05, only edges with lower p-values were selected as features. Vectors of edge values across subjects were created for the features surviving the threshold. These were then z-scored and input into PCA such that the PCs were coefficients for linear combinations of features. From the feature PCA for each iteration, we calculated the fraction of variance explained by each PC. Since these fractions sum to 1 and are non-negative, we treated them as a probability distribution and used them to calculate the normalized Shannon entropy. Each point represents the entropy value with the violin plots showing the distributions. This process was done for phenotypes age **(A)**, IQ **(B)**, and sex **(C)**.

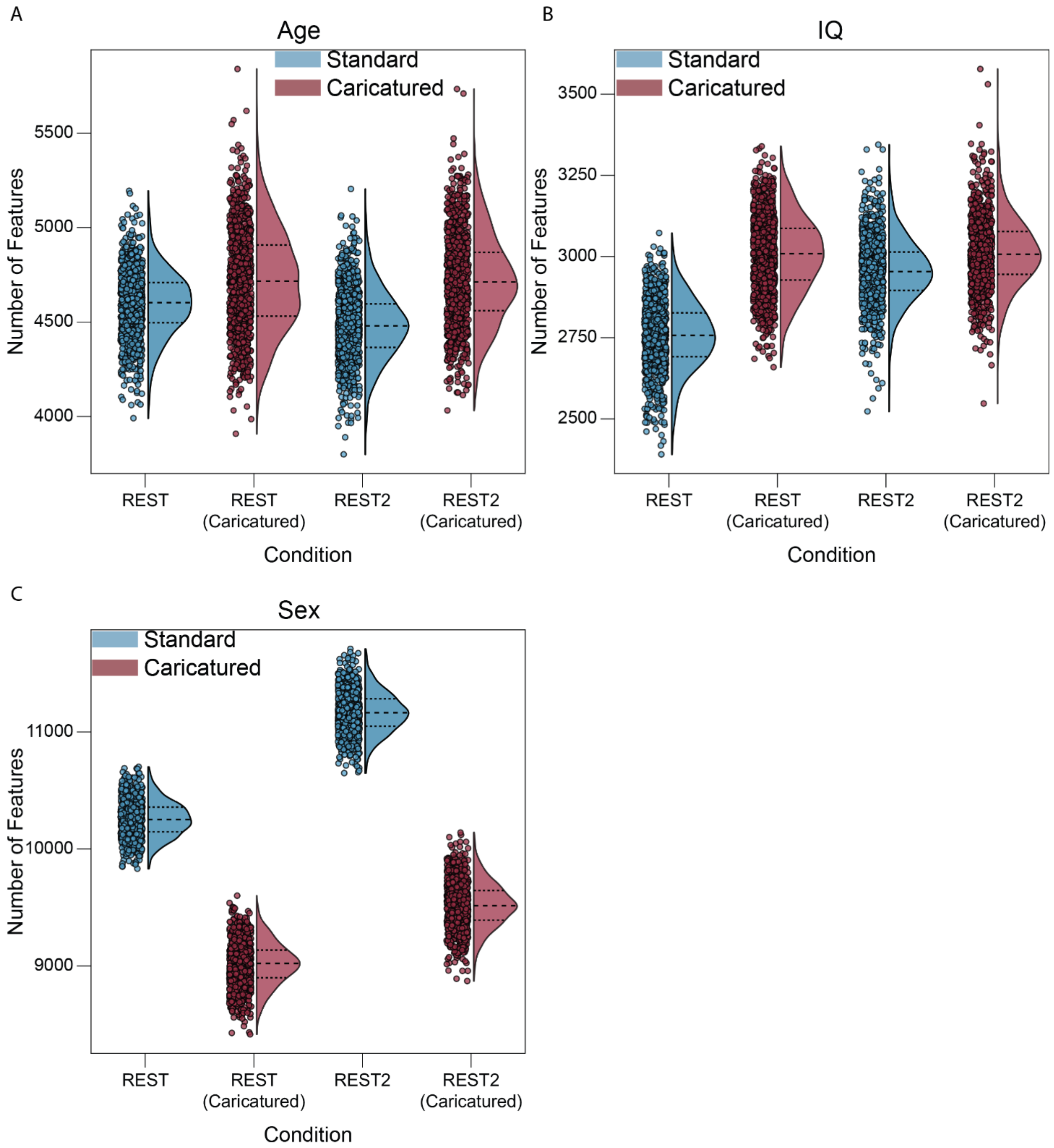


**SI Figure 8:** Number of features in each model. To estimate the features used in each of the 1000 models built in the CPM process, we iterated through each of the subsamples and found the p-value for the association between the phenotype and each edge across subjects. As the p-value threshold for the models was 0.05, only edges with lower p-values were selected as features. Here, the number of features that survived the threshold in each iteration were noted by the points on the graph, with the violin plot showing the distribution. This process was done for phenotypes age **(A)**, IQ **(B)**, and sex **(C)**.

**References**


1. Shine, J. M. *et al.* Human cognition involves the dynamic integration of neural activity and neuromodulatory systems. *Nature neuroscience* **22**, 289–296 (2019).
2. Finn, E. S. *et al.* Functional connectome fingerprinting: identifying individuals using patterns of brain connectivity. *Nat Neurosci* **18**, 1664–1671 (2015).
3. Noble, S. *et al.* Influences on the test–retest reliability of functional connectivity MRI and its relationship with behavioral utility. *Cerebral cortex* **27**, 5415–5429 (2017).
